# Characterization of NPR-14 in the Regulation of Sleep-Like Behaviour in *Caenorhabditis elegans*

**DOI:** 10.64898/2026.07.21.739920

**Authors:** Foroozan Torki, William G. Bendena, Ian D. Chin-Sang

## Abstract

Sleep-like quiescence is an evolutionarily conserved state essential for physiological homeostasis; however, its dysregulation can lead to sleep disorders such as narcolepsy, which can be caused by abnormal neuropeptide signaling. In *Caenorhabditis elegans*, the G-protein-coupled receptor NPR-14 belongs to the orexin/allatotropin receptor family and has been proposed as a homolog of mammalian orexin receptors. Using *npr-14* loss-of-function (*lf*) mutants, we demonstrate that NPR-14 promotes arousal and inhibits sleep-like quiescence. *npr-14(lf)* mutants exhibit prolonged quiescence, reduced locomotion, impaired sensory responses, and metabolic defects including elevated fat accumulation and decreased feeding and egg-laying. NPR-14 is expressed in ASH and ASI sensory neurons and in GABAergic DD, VD, and VC motor neurons, positioning it to modulate both sensory-motor integration and motor output directly. Genetic epistasis analysis revealed that NPR-14 functions upstream of EGL-4/protein kinase G (PKG): *egl-4* loss-of-function suppressed the enhanced quiescence of *npr-14* mutants, while *egl-4* gain-of-function phenotypes were not enhanced by loss of *npr-14*. Caffeine treatment partially suppressed *npr-14* mutant quiescence, suggesting convergence with adenosine-sensitive arousal pathways. These findings establish NPR-14 as a wake-promoting GPCR that inhibits EGL-4/PKG signalling to regulate quiescence and arousal. The NPR-14–EGL-4 axis suggests functional parallels to arousal regulation in other systems.

## Introduction

Sleep is a fundamental, evolutionarily conserved behaviour essential for cognitive performance, metabolic balance (1,2), and overall health. Even mild sleep loss impairs attention, memory (3), and decision-making, while chronic disruption increases the risk of cardiovascular disease (4,5), metabolic disorders, immune dysfunction (6), and neuropsychiatric conditions. In humans, sleep–wake regulation depends on complex interactions between circadian and homeostatic processes (7).

Wakefulness is maintained by neurotransmitters such as dopamine, norepinephrine, histamine, and orexin, which promote cortical and thalamic activation (8–10). In contrast, sleep initiation arises from coordinated activity within sleep-promoting networks through specific neuropeptidergic signaling (11,12). Sleep-promoting systems also include GABA and galanin from the ventrolateral preoptic nucleus, which silence wake-promoting nuclei (13,14), and somatostatin, which contributes to sleep regulation (14–16). The interplay between these systems regulates the transitions between wakefulness, and sleep; accordingly, dysregulation of this signaling is a key driver in various sleep disorders (9,17),(18).

Orexin (hypocretin), produced in the lateral hypothalamus, stabilises wakefulness by exciting monoaminergic systems via two GPCRs, OX1R and OX2R (8,9,19). Loss of orexin signalling results in narcolepsy with cataplexy (20). Counterbalancing this wake drive is adenosine, which accumulates with prolonged wakefulness and promotes sleep via A1 receptors (19,21–23). Caffeine antagonises adenosine receptors, highlighting the biological importance of this pathway (24).

Studies in *Drosophila melanogaster* have identified conserved sleep-regulatory mechanisms. Fly sleep is defined behaviourally as periods of immobility exceeding five minutes, accompanied by increased arousal thresholds and homeostatic rebound (25,26). The allatostatin-A receptor DAR-2, which phylogenetically belongs to the galanin receptor family, promotes sleep by dampening excitatory drive and decreasing insulin release (27–30). Notably, the allatotropin receptor—structurally related to vertebrate orexin receptors—has been lost in *Drosophila* (31,32), raising the possibility that different GPCR families may have evolved opposing roles in sleep regulation.

cGMP-dependent protein kinase (PKG) represents a conserved downstream effector of sleep regulation. In *Drosophila*, the *foraging* (*for*) gene encodes PKG; high-PKG “rover” strains show greater resilience to sleep deprivation than low-PKG “sitter” strains (33). While PKG promotes sleep in *C. elegans*, high PKG levels in *Drosophila* increase wakefulness, suggesting opposite behavioural outputs despite molecular conservation (33). In mammals, PKG similarly promotes sleep and opposes wake-promoting cAMP/PKA pathways.

In *C. elegans*, sleep-like states occur as developmentally timed sleep (DTS) during lethargus (34,35) and stress-induced sleep (SIS) in response to heat shock, injury, or infection (35–37). These states exhibit conserved hallmarks including reduced locomotion, diminished pharyngeal pumping, increased arousal thresholds, and homeostatic rebound (38). SIS induction requires EGF signalling through LET-23/EGFR, which activates the sleep-promoting neurons ALA and RIS (36,39,40). ALA releases FLP-13 neuropeptides that act via DMSR-1 (41–43), while RIS releases FLP-11 neuropeptides (44,45). These inhibitory neuropeptides suppress cAMP/PKA signalling via Gi/o-coupled receptors including DMSR-1 and FRPR-4 (46,47).

Opposing this pathway, EGL-4 (PKG) serves as a major sleep-promoting effector. EGL-4 functions in sensory neurons to modulate arousal thresholds and locomotion (34,46,48); gain-of-function mutations cause animals to slow or stop, while loss-of-function causes hyperactivity and reduced sleep. EGL-4 promotes sleep by antagonising cAMP-dependent signalling, establishing a conserved regulatory axis that determines neuronal excitability.

NPR-14 is a *C. elegans* G-protein-coupled receptor belonging to the orexin/allatotropin receptor family (49). Phylogenetic analyses place NPR-14 within this conserved family that includes vertebrate orexin receptors (OX1R, OX2R) and invertebrate allatotropin receptors, (Figure 1). These receptors share conserved structural features and are all implicated in the regulation of arousal and metabolism. Given the established roles of orexin receptors in vertebrate arousal, we investigated whether NPR-14 functions as a sleep-regulatory GPCR in *C. elegans*, examining its expression patterns, effects on behaviour and metabolism, and genetic interaction with the EGL-4/PKG pathway. Our findings reveal that NPR-14 is a wake-promoting GPCR whose inhibition of EGL-4/PKG signaling plays a critical role in modulating transitions between quiescence and arousal.

**Figure 1.**
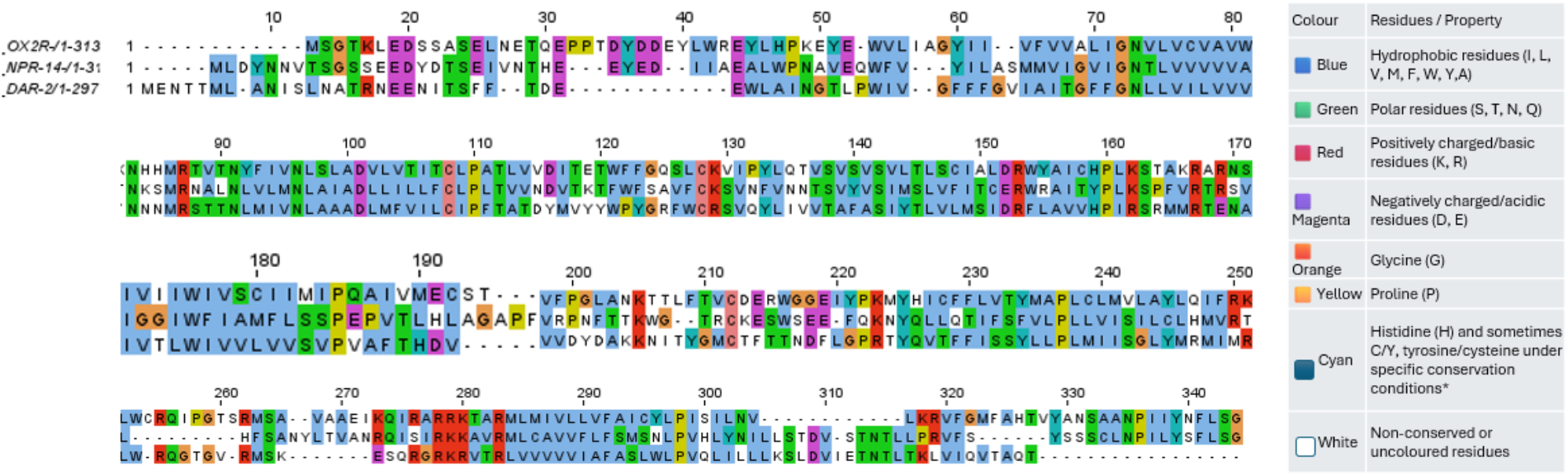
Amino acid sequence alignment of *Caenorhabditis elegans* NPR-14, *Drosophila melanogaster* DAR-2, and human orexin receptor 2 (OX2R). The alignment was generated using Jalview (50), with conserved residues colored according to physicochemical similarity. Pairwise sequence comparisons show that NPR-14 and OX2R share 53% similarity and 34.66% identity, while NPR-14 and DAR-2 share 47.2% similarity and 25.1% identity.

## Materials and methods

### *C. elegans* maintenance

The nematodes were cultured on nematode growth medium (NGM), a standard agar-based medium widely used for *C. elegans* cultivation at a temperature of 20°C., within the optimal range for their growth and reproduction*. E.coli* OP50, a non-pathogenic strain, was pipetted onto the NGM plates to serve as a food source for the worms (64). A list of strains used in this study is presented in (Table 1). Because the behavioural and physiological phenotypes of both *npr-14(ok2375lf)* and *npr-14(tm2974lf)* alleles were highly similar across assays, we present the results from both alleles for completeness (**S2**). However, because *ok2375* carries the larger deletion, the data presented here are based primarily on *npr-14(ok2375lf)* as the principal mutant allele.

**Table 1:** *C. elegans* Strains used in this research.

| Strain | Genotype | Origin |
| --- | --- | --- |
| N2 | Wild-type | CGC |
| RB1836 | *npr-14(ok2375lf)* | CGC |
| IC1116 | *npr-14(tm2974lf)* | CGC |
| AD521 | *egl-4(ad450gf)* IV | CGC |
| MT1074 | *egl-4(n479lf)* IV | CGC |
| IC807 | *quEx212 [npr-14p::mCherry]* | Queen's University, Chin-Sang Laboratory |
| IC2631 | *npr-14(ok2375lf); egl-4(ad450gf)* | Queen's University, Chin-Sang Laboratory |
| IC1117 | *npr-14(ok2375lf); egl-4(n479lf)* | Queen's University, Chin-Sang Laboratory |
| IC870 | *quEx239[npr-14(+)]* | Queen's University, Chin-Sang Laboratory |
| IC871 | *quEx240[npr-14(+)]* | Queen's University, Chin-Sang Laboratory |
| IC874 | *quEx242[npr-14(+)]* | Queen's University, Chin-Sang Laboratory |
| IC875 | *quEx243[npr-14(+)]* | Queen's University, Chin-Sang Laboratory |
| IC876 | *quEx244[npr-14(+)]* | Queen's University, Chin-Sang Laboratory |
| IC2609 | *npr-14(ok2375lf); quEx244[npr-14(+)]* | Queen's University, Chin-Sang Laboratory |

### *npr-14(+)* rescue

To generate *npr-14(+)* transgenic worms, 9 kbp of *npr-14* genomic DNA was amplified by PCR using primers (Table 2). 30ng/μl *C. elegans* PCR product along with 30ng/μl *Psur5::gfp*, injected directly into *npr-14(ok2375lf)* and *npr-14(tm2974lf)* mutants worms following standard *C. elegans* microinjection techniques (65). The injected nematodes were then allowed to recover and reproduce. Five independent *npr-14 (+)* rescue lines*, quEx239-quEx244*, were isolated (see Table 1). Rescue experiments were performed using the extrachromosomal array carrying *npr-14(+)* genomic DNA *quEx244[npr-14(+)]*.

**Table 2:**
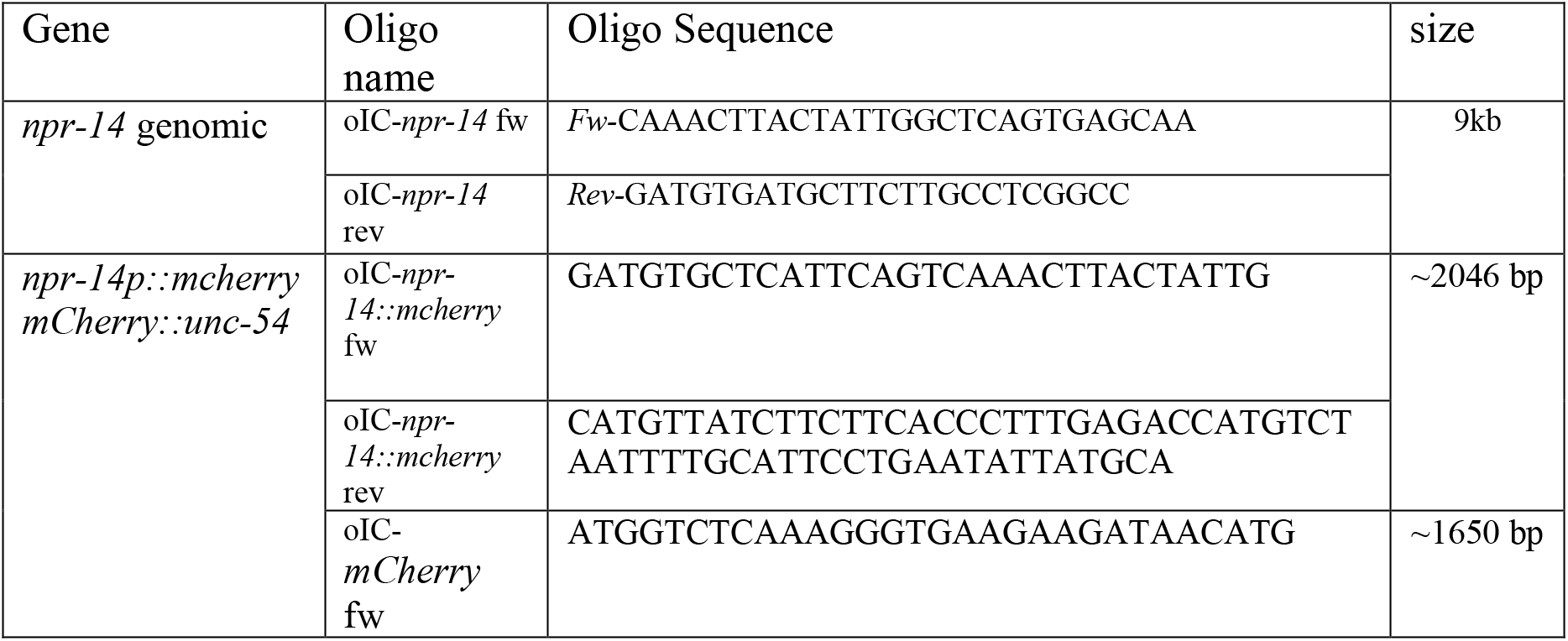
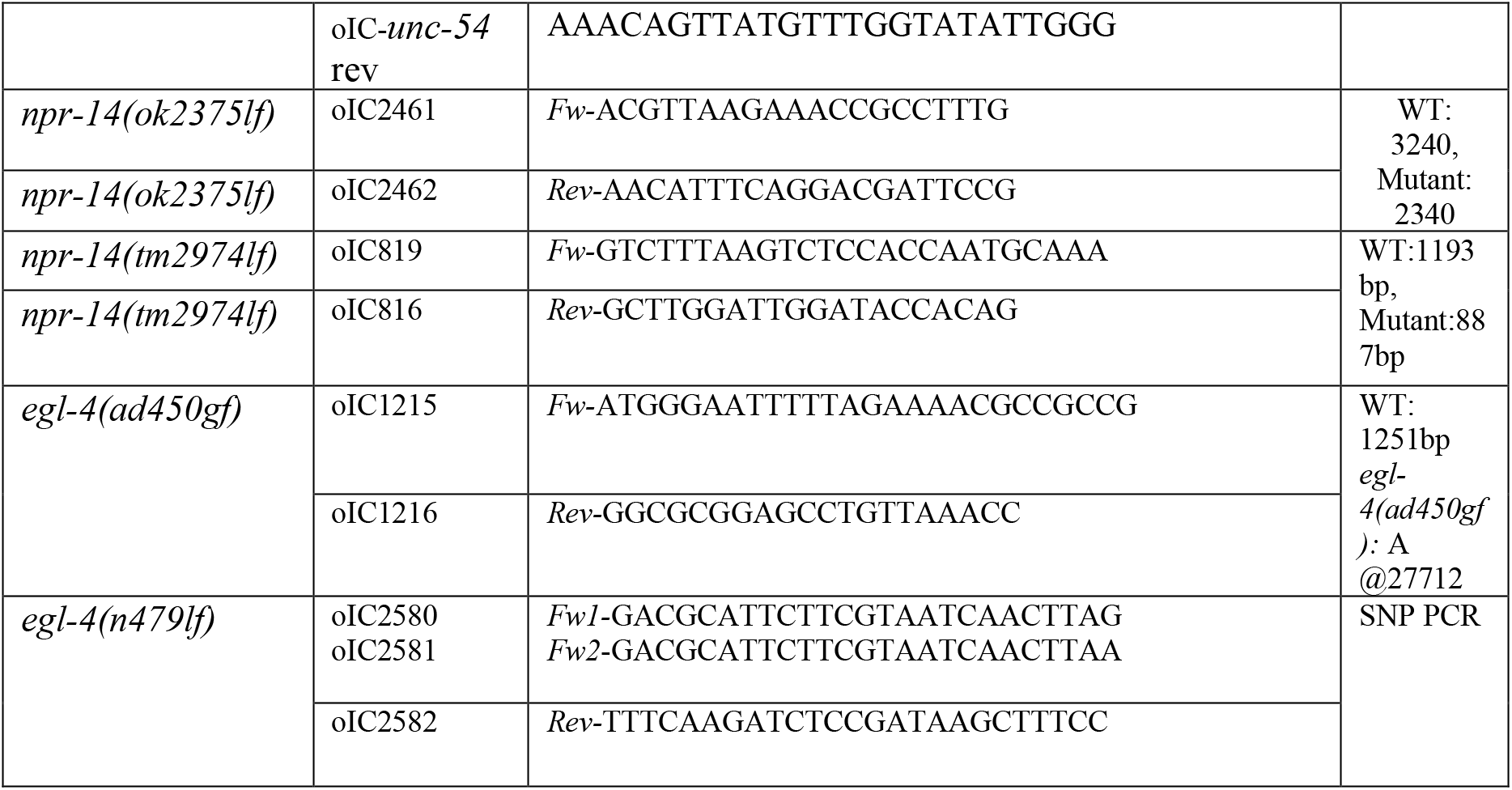
Oligo sequences.

### Generation of the *npr-14(ok2375lf); egl-4(ad450gf)* and *npr-14(ok2375lf); egl-4(n479lf)* double mutants

To obtain *npr-14(ok2375lf) I; egl-4(ad450gf)IV* double mutants, we performed marker-assisted genetic crosses. Males of the genotype *zdIs5[mechanosensory neuron GFP] I; mIs11[pharyngeal GFP] IV; him-5(e1490) V* were crossed with *npr-14(ok2375lf)* hermaphrodites. F₁ progeny expressing both GFP markers were heterozygous for *npr-14(ok2375lf)* and subsequently crossed with *egl-4(ad450gf)* homozygotes. From this cross, animals carrying non-GFP mechanosensory neuron and GFP pharyngeal (*+/ok2375; mIs11/ad450; him-5/+*) were allowed to self-fertilize. Non-fluorescent F₂ progeny were picked. Double mutants *npr-14(ok2375lf); egl-4(ad450gf)* were confirmed by PCR genotyping (see Table 2) for a list of primers and sequences. This marker-assisted approach was also used to make the *npr-14(ok2375lf); egl-4(n479lf)* double mutants.

### *npr-14p::mCherry* Reporter

The *npr-14p::mCherry* transcriptional reporter construct was generated by stitching PCR (66). A 2,016 bp upstream regulatory region of *npr-14* was amplified from N2 genomic DNA using the forward primer *npr-14p* and a reverse primer containing a 30 bp overlap with the 5′ end of *mCherry*. The *mCherry::unc-54 3′UTR* cassette was amplified from pCFJ90-pmyo-2::*mCherry::unc-54* 3′UTR using an *mCherry* forward primer and an *unc-54* 3′UTR reverse primer. The two purified PCR products were then fused by overlap-extension PCR using the forward *npr-14p* and reverse *unc-54* primers to generate the final *npr-14p::mCherry::unc-54 3′UTR* reporter construct.

### FITC (Fluorescein Isothiocyanate) Staining

To confirm the identity of neurons expressing *npr-14::mCherry*, FITC staining was used to label the amphid sensory neurons ASH, ASI, ASJ, ASK, ADL, and ADF (67). *npr-14::mCherry* worms were washed with M9 solution and incubated in 1 mg/ml FITC in M9 at 20 °C for 2 hours. Worms were then transferred to seeded NGM plates and allowed to recover for at least 2 hours to excrete residual dye. After 2–3 hours, co-localization of FITC and *npr-14::mCherry* fluorescence was examined.

### Roaming assay

Roaming assays were conducted as previously described (62). Briefly, synchronized early adults were transferred to seeded plates and acclimatized at 20°C for 60 minutes before movement behaviour was assessed. Roaming/searching behaviour was considered positive if the animal travelled a distance greater than four times its body length within 5 minutes. If the animals remained inside a perimeter roughly four times their body length, they were classified as pivoting/local search behaviour. Each experiment had a minimum sample size of 30 animals.

### Fat accumulation assay

Lipid content was assessed by cultivating synchronized nematodes to the adult stage, followed by staining with Oil-Red-O (ORO) in accordance with established staining and quantification protocols (68,69). ORO, a lipophilic dye employed for fatty tissue staining in *C. elegans*, selectively binds to lipids, facilitating the visual observation and quantitative measurement of accumulated fat within the nematodes. Worms were fixed in a solution containing 1× MRWB buffer supplemented with 1% paraformaldehyde. Fixation was performed by gentle rocking at room temperature for 1 hour. Fixed worms were washed with 1× PBS, resuspended in 60% isopropanol, and incubated at room temperature for 15 minutes. Staining was performed using a saturated ORO solution dissolved in 60% isopropanol overnight with continuous rocking. Following staining, worms were washed twice with 1× PBS containing 0.01% Triton X-100 to remove excess dye (68). The imaging experiments were conducted using the ZEN 2010 software suite (Carl Zeiss MicroImaging GmbH) using brightfield and laser-scanning confocal microscopy. The system was configured with a zoom factor of 0.6, a master gain of 721, a digital gain of 1.00, and a pinhole aperture of 80 µm. Fluorescence detection employed a band-pass filter spanning 548–690 nm, and excitation was achieved using a 543 nm laser line. Relative fat accumulation was determined by comparing mutant relative fluorescence (RF) intensities to those of wild-type worms.

### Pharyngeal pumping assay

This method was adapted from (37). To assess feeding behaviour, 10–15 adult worms were transferred to NGM plates seeded with *E. coli* OP50 and allowed to acclimate for 15 minutes prior to the assay. Feeding rate was measured by observing the rhythmic contractions of the grinder located in the terminal bulb of the pharynx. Each complete up-and-down movement was counted as one pharyngeal pump. The number of pumps was recorded over a 10-second interval for each worm using a dissecting microscope.

### Head Thrashing Assay

To assess locomotor activity and disruption of quiescence, we performed a thrashing assay in a liquid medium under both control (no caffeine) and caffeine-treated conditions. Adult *C. elegans* (10–15 per group) were transferred into a 30 µL droplet of solution (M9 buffer or M9 containing 15mM caffeine) placed on a glass slide (with a coverslip “bridge” to prevent compression). After a 5-minute acclimation period, the number of worms that continued thrashing was recorded, and thrashing frequency was quantified per 10 seconds with one full lateral body bend (one complete swing) defined as a single thrash (Figure 8) or every 5 minutes for 20 min (i.e., at 5, 10, 15, and 20 min) and at each time point, thrashing frequency was quantified for 30 seconds per worm (Figures 10,11). For caffeine treatment, we selected 15 mM caffeine, a dose previously used in *C. elegans* locomotion studies (68,69). We compared temporal changes between control and caffeine conditions across genotypes.

### Egg-Laying Assay

This method was adapted from previously described protocols for quantifying *C. elegans* egg-laying behaviour (69). Egg-laying behaviour was quantified to assess reproductive output in *C. elegans*. L4-stage hermaphrodites were isolated and transferred individually to separate NGM plates seeded with *E. coli* OP50. Worms were maintained at 20 °C, and the number of eggs laid by each animal was counted every 24 hours. After each count, adults were transferred to fresh seeded plates to prevent confusion with previously laid eggs. The assay continued until the worms ceased egg production. For each strain, 10–15 worms were analyzed per biological replicate, and three independent replicates were performed. The mean number of eggs laid per worm per day was calculated to determine the temporal pattern and total reproductive output.

### Octanol response assay

To assess aversive response to chemical stimuli, 10–15 adult *C. elegans* were transferred to an unseeded NGM plate, ensuring that no *E. coli* OP50 bacteria were carried over during transfer. Worms were allowed to acclimate on the plate for 15 minutes prior to testing. For the assay, a fine paintbrush bristle was dipped in 30% 1-octanol and gently positioned near the worm’s head, without making physical contact. A behavioural response is defined as either a backward movement or head withdrawal, and the time to respond was noted for each worm (37).

### Quiescence assay

Adult worms (5–6 per assay) were transferred to NGM plates containing 25 μL of *E. coli* OP50. After transfer, animals were allowed to acclimate for 15 minutes at room temperature. Following acclimation, worm behaviour was recorded. Video data were processed and analyzed using the WormLab Imaging System (MBF Bioscience).

### Statistical analysis

All data visualization and statistical analyses were performed using GraphPad Prism version 9.4.1 (GraphPad Software, San Diego, California, USA). Behavioural data from the quiescence and pharyngeal pumping assays were analyzed using one- and two-way ANOVAs to assess differences. Post-hoc comparisons were conducted using Tukey–Kramer multiple comparisons tests to determine statistically significant differences between groups.

For the 1-octanol response assay, differences between groups were analyzed using Fisher’s exact test, following Hill et al. (2014)(35). A p-value of <0.05 was considered statistically significant for all tests.

## Results

### Neuronal Localization of *npr-14*

To determine the expression pattern of *npr-14*, we generated a transcriptional reporter construct (*npr-14p::mCherry*) and visualized *mCherry* fluorescence using confocal microscopy. This analysis revealed robust expression in the ASH and ASI sensory neurons (Figure 2). Both ASH and ASI are amphid neurons extend dendritic processes to the external environment through openings in the cuticle. The amphid identity of these neurons was confirmed by co-labelling with FITC, a fluorescent dye selectively taken up by amphid sensory neurons (see Methods). Reporter expression was also detected in the body and tail regions. In the mid-body, *npr-14p::mCherry* fluorescence was observed in VC4 near the vulva. Additional expression was detected in VD and DD motor neurons along the ventral nerve cord, including commissural processes connecting the ventral and dorsal cords. In the tail, expression was consistent with VD13 and DD6, although additional *npr-14*-expressing neurons were also present (not shown).

**Figure 2.**
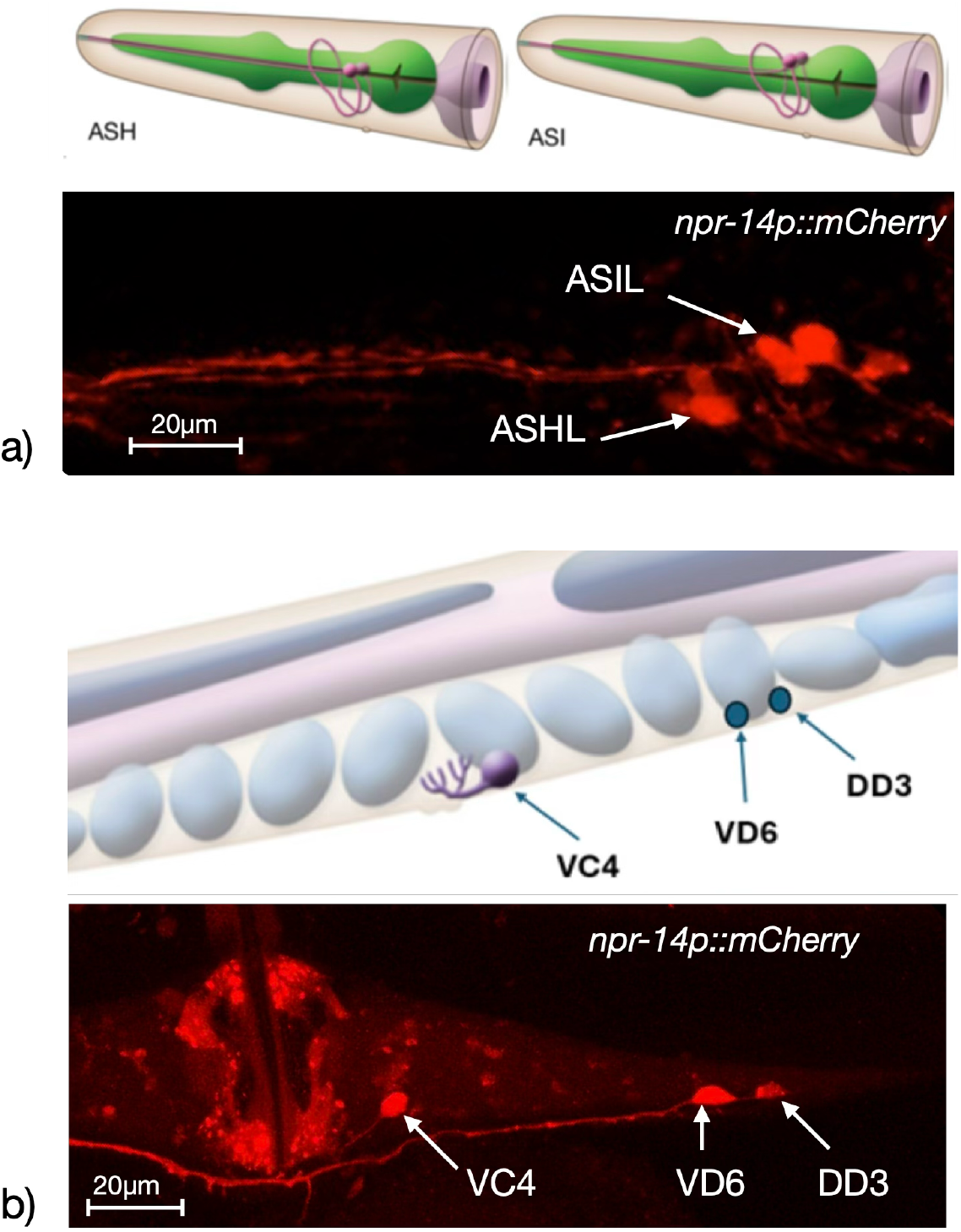
*npr-14p::mCherry* expression in sensory and motor neurons. (a) WormAtlas-based schematic representations of ASH and ASI amphid sensory neurons, together with a confocal image showing *npr-14* promoter-driven *mCherry* expression in the anterior head region. Fluorescent cell bodies and dendritic processes are observed in positions consistent with ASH and ASI sensory neurons, including ASHL and ASIL. (b) WormAtlas-based schematic representation of ventral cord neurons, including VC, VD, and DD neurons. The corresponding confocal image shows *npr-14p::mCherry* expression along the ventral nerve cord, with labelled neuronal cell bodies consistent with VC4, VD6, and DD3. Fluorescent signal is also visible in neuronal processes extending along the body axis.

### Behavioural Assays

To test whether *npr-14* regulates sleep-like quiescence, we examined behavioural assays previously established as markers of sleep states in *C. elegans* (34). Sleep-like quiescence in worms is characterized by decreased locomotor activity, reduced feeding, and diminished responsiveness to external stimuli. We therefore measured: (i) roaming behaviour to assess spontaneous locomotor activity; (ii) octanol avoidance to test responsiveness to aversive chemical stimuli; and (iii) pharyngeal pumping rate to evaluate feeding behaviour.

### NPR-14 Loss-of-Function Reduces Roaming Behaviour

*npr-14(ok2375lf)* loss-of-function mutants exhibited significantly reduced roaming compared to wild-type animals (Figure 3). This defect was rescued by transgenic expression of a wild-type *npr-14* genomic construct (*quEx244[npr-14(+)]*), confirming that the observed phenotype results from loss of NPR-14 function. These behavioural findings are consistent with NPR-14 expression in ASH and ASI sensory neurons, which regulate foraging and arousal (51) and in DD and VD GABAergic motor neurons, which directly control locomotion (52).

**Figure 3.**
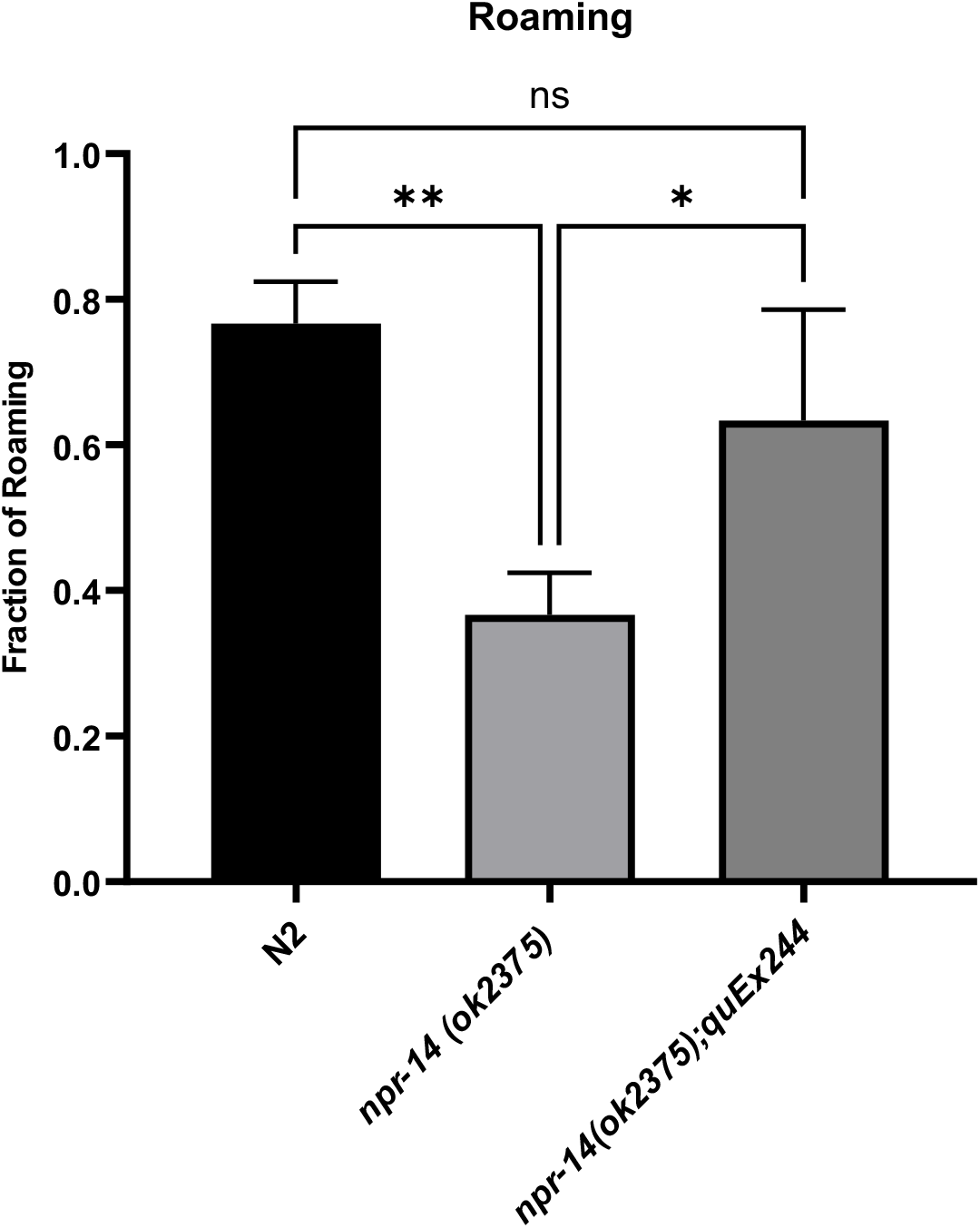
NPR-14 loss-of-function reduces roaming behaviour. Quantification of roaming behaviour in wild-type N2, *npr-14(ok2375lf)* mutants, and rescued *npr-14(ok2375lf);quEx244[npr-14(+)]* animals. *npr-14(ok2375lf)* mutants show significantly reduced roaming compared to wild-type. Transgenic expression of wild-type *npr-14* restores roaming to wild-type levels. Data are mean ± SEM; *p < 0.01, **p < 0.001, ns = not significant.

### NPR-14 Loss-of-Function Reduces Pharyngeal Pumping

*npr-14(ok2375lf)* mutants displayed significantly reduced pharyngeal pumping rates compared to wild-type N2 animals (Figure 4). This reduction in pumping frequency indicates decreased feeding activity. Transgenic expression of wild-type *npr-14 quEx244[npr-14(+)]*restored pumping rates to wild-type levels, confirming that the feeding defect results from loss of NPR-14 function.

**Figure 4.**
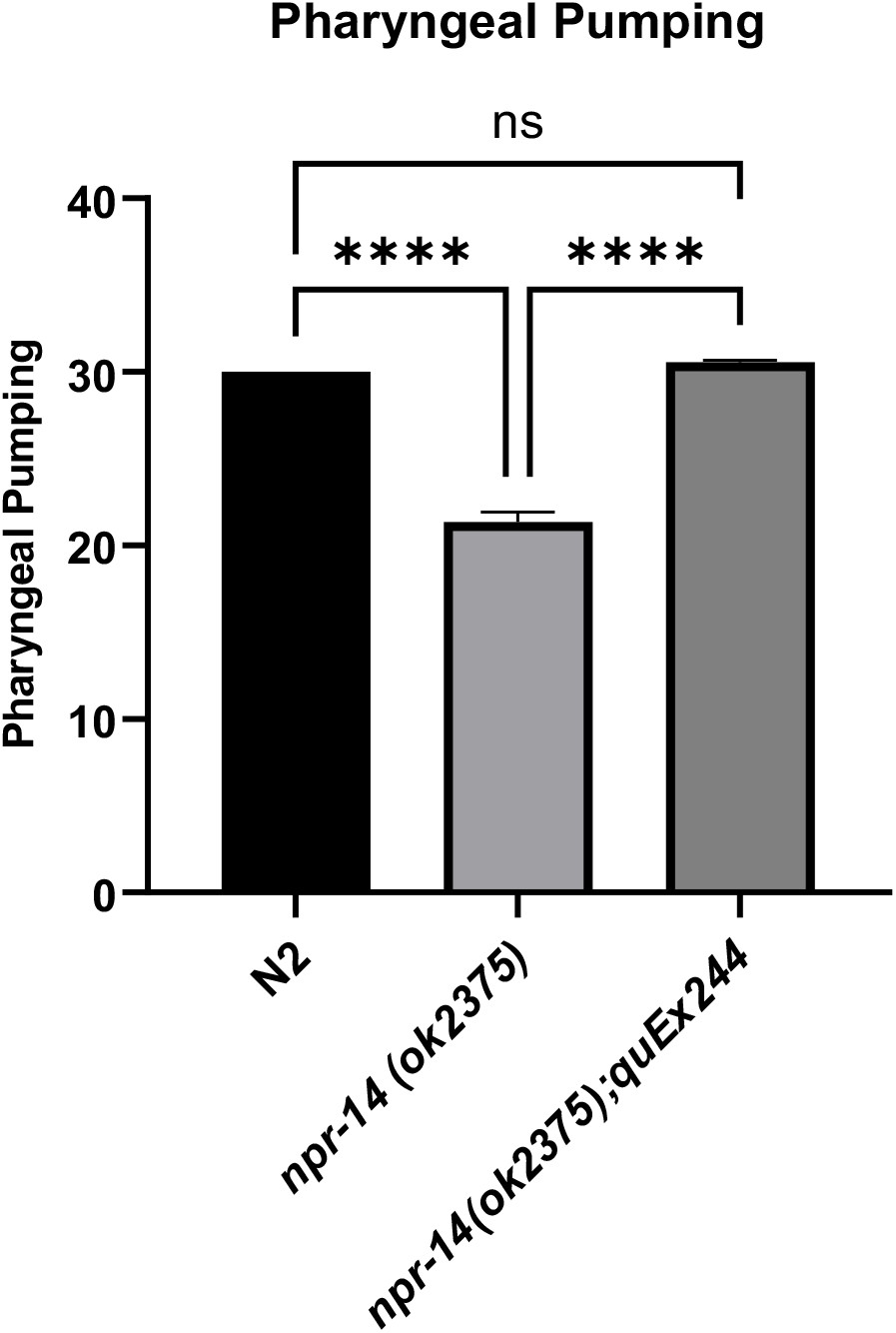
NPR-14 loss-of-function reduces pharyngeal pumping. Pharyngeal pumping rates in wild-type N2, *npr-14(ok2375lf)* mutants, and rescued *npr-14(ok2375);quEx244[npr-14(+)]* animals. *npr-14(ok2375lf)* mutants exhibit significantly reduced pumping compared to wild-type. Transgenic expression of wild-type *npr-14* restores pumping to wild-type levels. Data are mean ± SEM (pharyngeal contractions per 10 seconds); ***p < 0.0001, ns = not significant.

### NPR-14 Loss-of-Function Impairs Sensory Responsiveness

To assess whether NPR-14 regulates arousal threshold, we measured avoidance latency to octanol. *npr-14(ok2375lf)* mutants exhibited significantly delayed responses to octanol compared to wild-type N2 animals (Figure 5). This defect was rescued by transgenic expression of wild-type *npr-14 quEx244[npr-14(+)]*, which restored avoidance responses to near wild-type levels. These results confirm that NPR-14 is required for normal sensory responsiveness.

**Figure 5.**
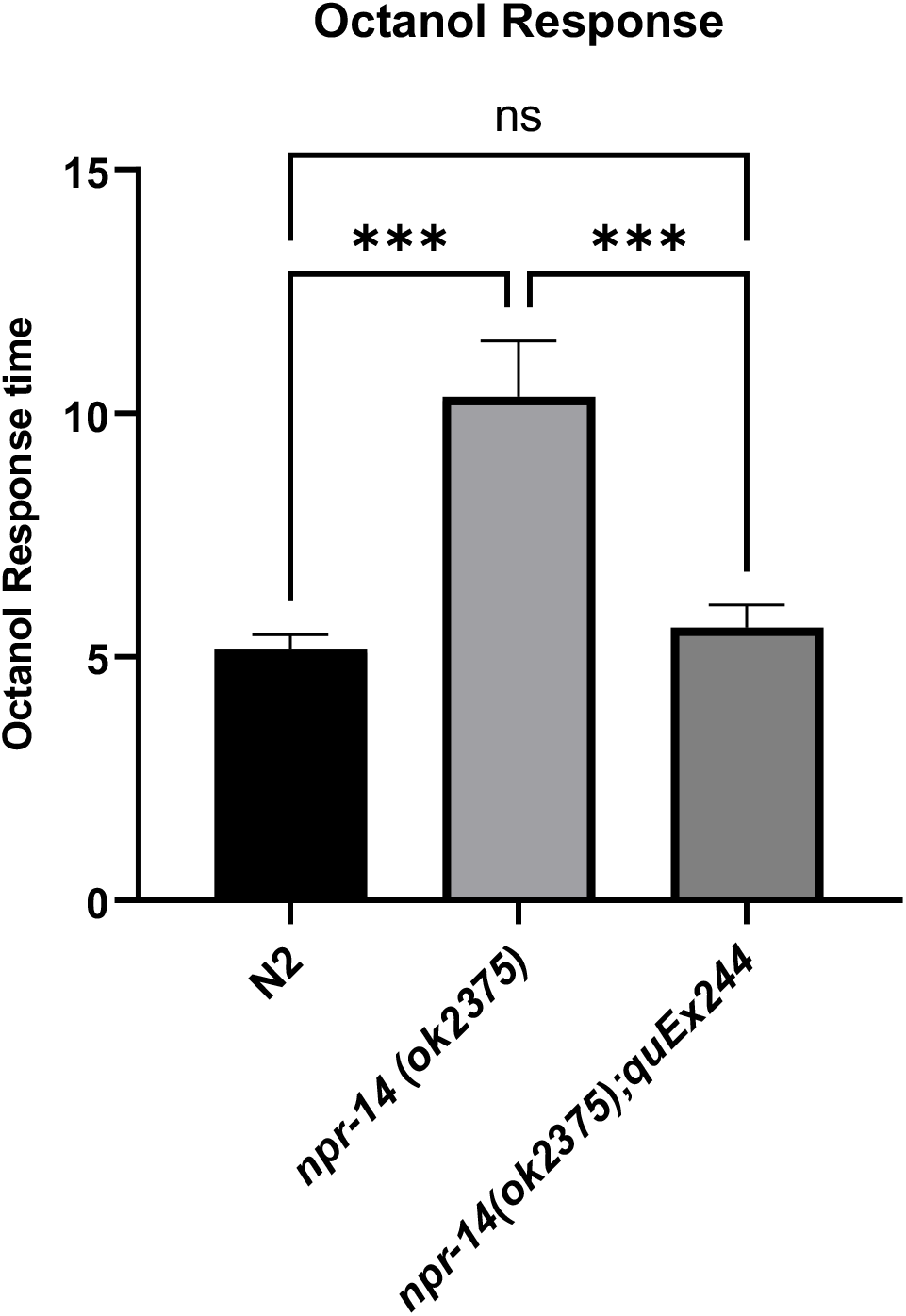
NPR-14 loss-of-function impairs octanol avoidance. Time (seconds) to respond to 30% octanol, wild-type N2, *npr-14(ok2375lf)* mutants, and rescued *npr-14(ok2375lf);quEx244[npr-14(+)]* animals. *npr-14(ok2375lf)* mutants exhibit significantly delayed responses compared to wild-type. Transgenic expression of wild-type *npr-14* restores responsiveness to near wild-type levels. Data represent mean ± SEM (seconds); ***p < 0.001, ns = not significant.

### NPR-14 Loss-of-Function Increases Lipid Storage

Sleep-like quiescence is closely coupled to metabolic regulation. We therefore examined whether *npr-14* loss-of-function affects energy homeostasis, reasoning that decreased locomotor activity and altered feeding behaviour in *npr-14* mutants may be accompanied by changes in lipid storage. We quantified fat accumulation using Oil Red O staining as a physiological readout of metabolic state. *npr-14(ok2375lf)* mutants displayed significantly elevated fat accumulation compared to wild-type N2 animals (Figure 6). This increase in lipid storage was rescued by transgenic expression of wild-type *npr-14 quEx244[npr-14(+)]*, which restored fat levels to those of wild-type animals.

**Figure 6.**
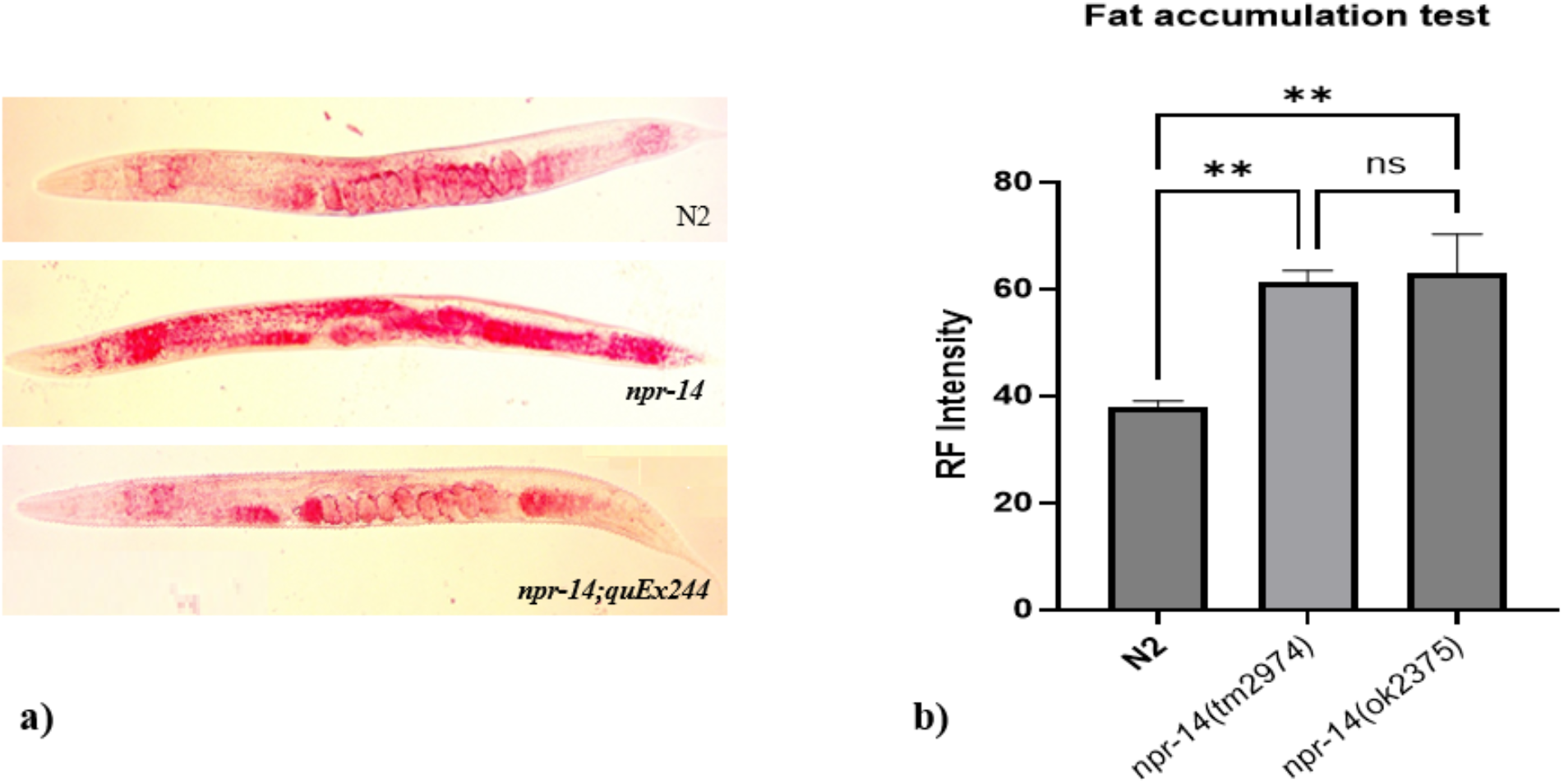
NPR-14 loss-of-function increases lipid accumulation. **(a)** Representative Oil Red O staining of lipid droplets in wild-type N2 and *npr-14(ok2375lf)* mutants. *npr-14(ok2375lf)* mutants show increased numbers and sizes of lipid droplets compared to N2. Transgenic rescue (*npr-14(ok2375);quEx244[npr-14(+)]*) restores lipid levels to wild-type. **(b)** Quantification of fat accumulation, Relative Fluorescence (RF). *npr-14(ok2375lf)* mutants exhibit significantly elevated fluorescence intensity compared to wild-type N2. Transgenic expression of wild-type *npr-14* restores fat accumulation to wild-type levels. Data are mean ± SEM; **p < 0.001, ***p < 0.0001, ns = not significant.

### NPR-14 Loss-of-Function Reduces Egg-Laying

In *C. elegans*, the cGMP-dependent protein kinase EGL-4 regulates behavioural state and has long been associated with both egg-laying control and quiescence-related phenotypes (53,54). Egg-laying alternates between active and inactive states and is coordinated with the animal’s overall motor state. Recent work has shown that egg-laying is reduced during stress-induced sleep, supporting the view that reproductive behaviour is suppressed during sleep-like quiescence (55). Reduced egg-laying in *npr-14* mutants is consistent with a general state of quiescence; however, we cannot exclude independent effects of NPR-14 on the egg-laying circuit (53). We quantified reproductive output by measuring the number of eggs laid over a six-day period. *npr-14(ok2375lf)* mutants produced significantly fewer eggs than wild-type N2 animals (Figure 7). This phenotype aligns with NPR-14 expression in VC motor neurons, which innervate the vulval muscles and regulate egg-laying (56,57). Transgenic expression of wild-type *npr-14 quEx244[npr-14(+)]* restored egg-laying rates to wild-type levels, confirming that the defect arises from loss of NPR-14 function.

**Figure 7.**
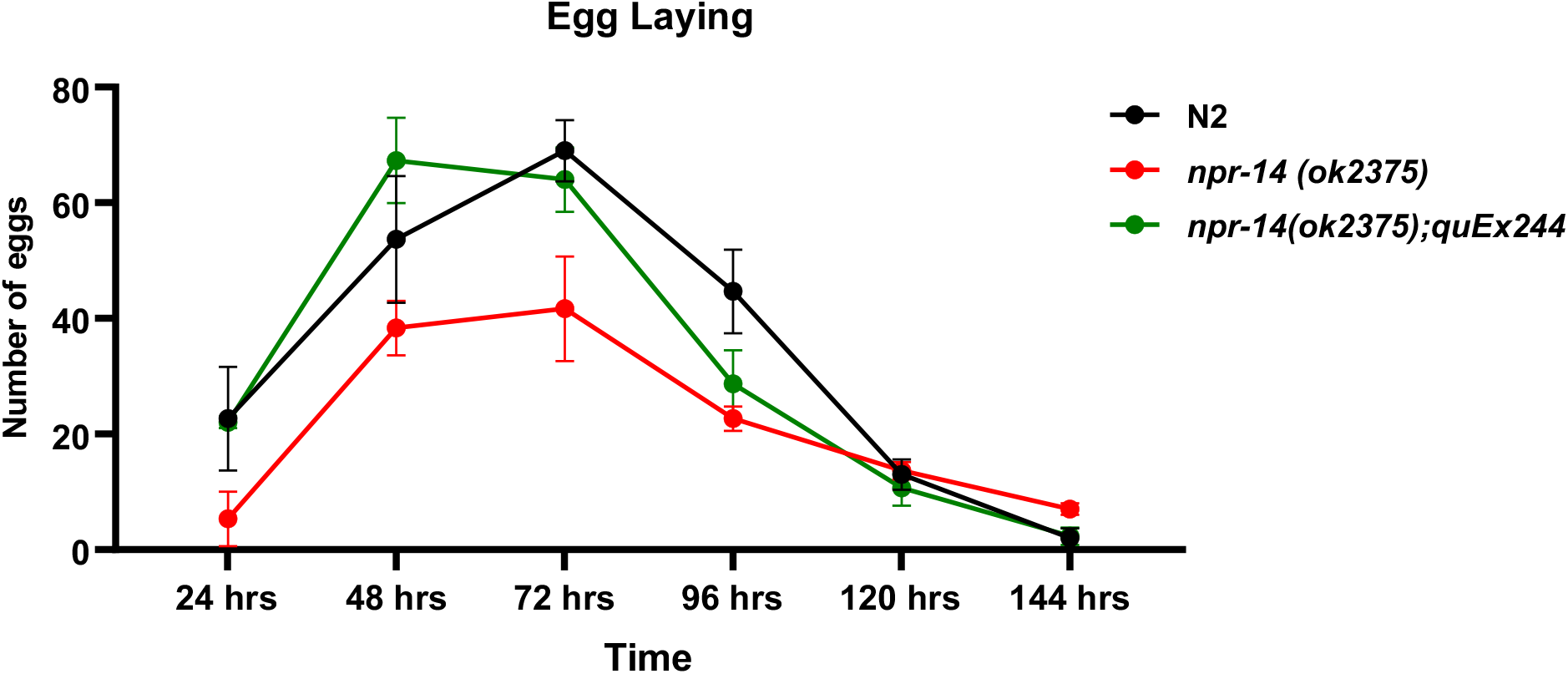
NPR-14 loss-of-function reduces egg-laying. Total eggs laid over six days by wild-type N2, *npr-14(ok2375lf)* mutants, and rescued *npr-14(ok2375);quEx244[npr-14(+)]* animals. *npr-14(ok2375lf)* mutants exhibit significantly reduced egg-laying compared to wild-type. Transgenic expression of wild-type *npr-14* restores egg-laying to wild-type levels. Data are mean ± SEM; statistical significance indicated.

### NPR-14 Loss-of-Function Reduces Thrashing Behaviour

Thrashing in liquid provides an additional quantitative measure of locomotor activity. Because quiescence in *C. elegans* is defined in part by reduced body movement, decreased thrashing serves as an independent indicator of lowered motor output consistent with sleep-like states. We quantified locomotor activity by measuring head thrashes per 10 seconds. *npr-14(ok2375lf)* mutants exhibited significantly reduced thrashing rates compared to wild-type N2 (Figure 8). Transgenic expression of wild-type *npr-14 quEx244[npr-14(+)]* restored thrashing to wild-type levels, confirming that the locomotor deficit results from loss of NPR-14 function. Notably, *npr-14(ok2375lf)* mutants often thrashed normally before suddenly stopping, a phenotype reminiscent of human narcolepsy (Supplementary Video S3).

**Figure 8.**
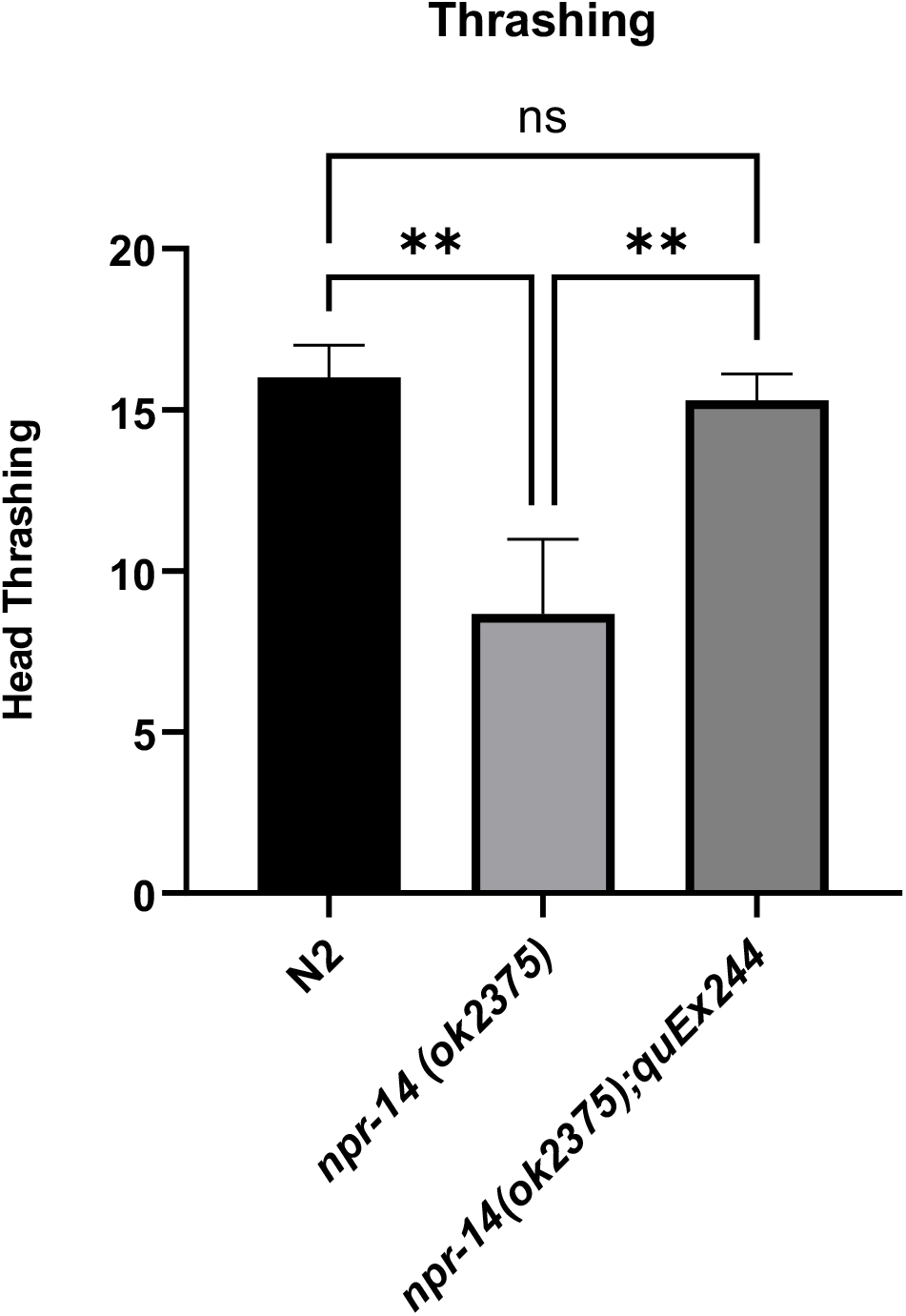
NPR-14 loss-of-function reduces thrashing behaviour. Head thrashes per 10 seconds in wild-type N2, *npr-14(ok2375lf)* mutants, and rescued *npr-14(ok2375);quEx244[npr-14(+)]* animals. *npr-14(ok2375lf)* mutants exhibit significantly reduced thrashing compared to wild-type. Transgenic expression of wild-type *npr-14* restores thrashing to wild-type levels. Data are mean ± SEM; *p < 0.05, **p < 0.01, ns = not significant.

### NPR-14 Loss-of-Function Increases Quiescence

Based on the behavioural and physiological phenotypes observed in *npr-14* mutants, we examined whether loss of *npr-14* directly affects sleep-like quiescence. Sleep in *C. elegans* was first formally defined by Raizen et al. (2008) (34), who demonstrated that lethargus-associated quiescence displays reversibility, reduced sensory responsiveness, and homeostatic regulation—criteria that distinguish sleep from inactivity. We adapted this framework to quantify quiescence in adult worms using automated movement tracking. *npr-14(ok2375lf)* mutants exhibited significantly increased quiescence duration compared to wild-type N2 (Figure 9), indicating that NPR-14 promotes wakefulness. Notably, *npr-14(ok2375lf)* animals often moved normally before suddenly becoming quiescent, a pattern reminiscent of narcoleptic episodes (Supplementary Video S4).

**Figure 9.**
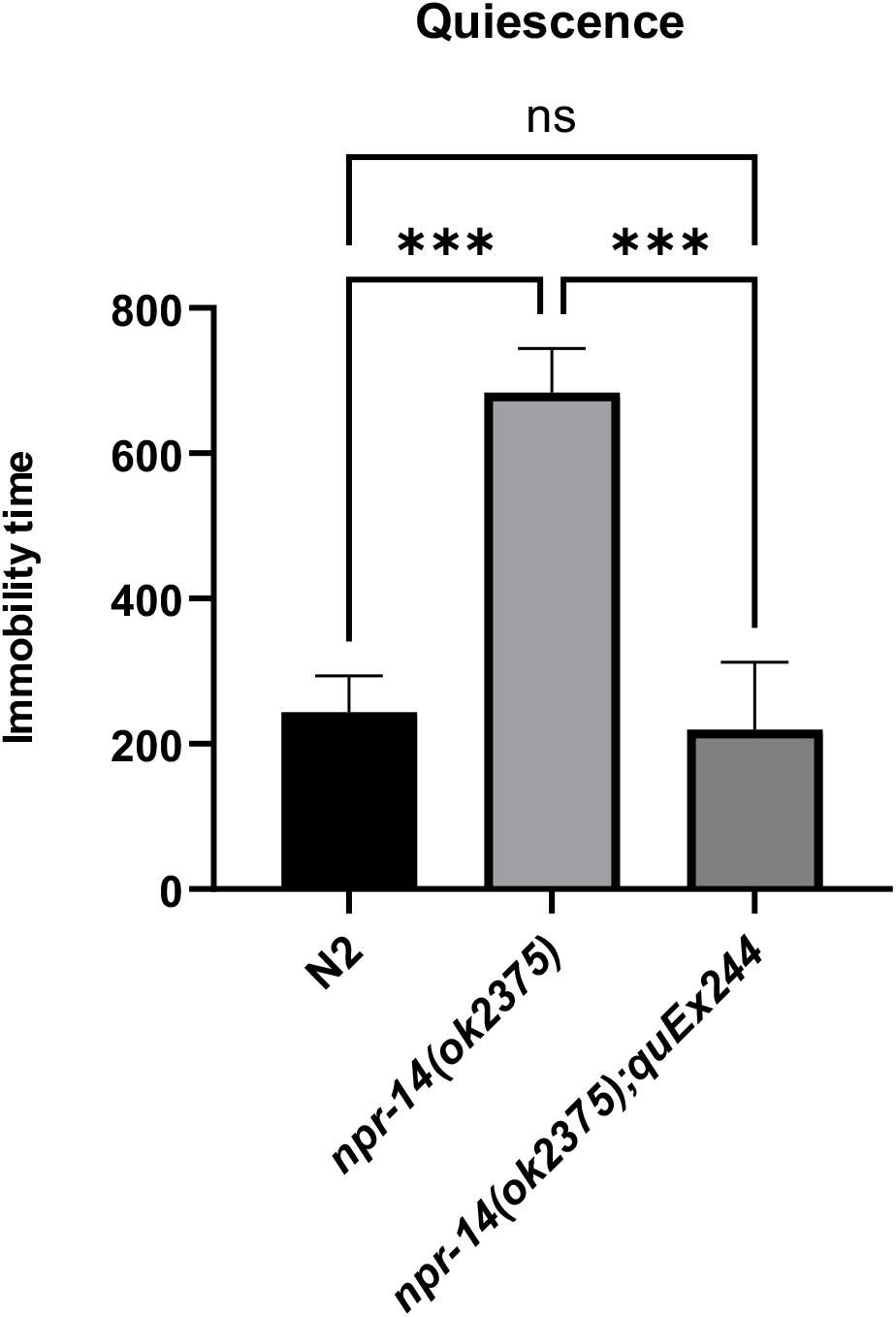
NPR-14 loss-of-function increases quiescence. Total quiescence duration (seconds) over a 30-minute period in wild-type N2, *npr-14(ok2375lf)* mutants, and rescued *npr-14(ok2375);quEx244[npr-14(+)]* animals. *npr-14(ok2375lf)* mutants exhibit significantly increased quiescence compared to wild-type. Transgenic expression of wild-type *npr-14* restores quiescence to wild-type levels. Data are mean ± SEM; ***p < 0.001, ns = not significant.

### NPR-14 Acts Upstream of EGL-4 to Inhibit Quiescence

To determine the genetic relationship between *npr-14* and the cGMP-dependent protein kinase *egl-4*, we performed epistasis analysis. EGL-4 is a well-characterized promoter of sleep-like quiescence: *egl-4(n479lf)* mutants exhibit reduced quiescence, while *egl-4(ad450gf)* mutants show increased quiescence (53). We first examined whether *npr-14* and *egl-4* function in the same genetic pathway. The *egl-4(ad450gf)* allele has quiescence to levels comparable to *npr-14(lf)* single mutants. Introducing the *npr-14(ok2375lf)* mutation into the *egl-4(ad450gf)* background did not significantly enhance quiescence beyond *egl-4(ad450gf)* alone (Figure 10a), indicating that *npr-14* and *egl-4* act in the same pathway rather than parallel pathways. To determine the epistatic relationship, we generated *npr-14(ok2375lf);egl-4(n479lf)* double mutants. The *egl-4(n479lf)* mutation completely suppressed the increased quiescence of *npr-14(ok2375lf)* mutants, with double mutants resembling *egl-4(n479lf)* single mutants (Figure 10a). This demonstrates that *egl-4* is epistatic to *npr-14*, placing *npr-14* upstream of *egl-4* in a linear pathway. Thus, NPR-14 normally acts to oppose or limit EGL-4-dependent quiescence to promote wakefulness; loss of NPR-14 leads to hyperactivation of EGL-4 and increased quiescence. We confirmed this epistatic relationship using the thrashing assay. *egl-4(n479lf)* mutants exhibited elevated thrashing (reduced quiescence), while *egl-4(ad450gf)* mutants showed reduced thrashing (increased quiescence). Consistent with the quiescence data, *npr-14(ok2375lf)* did not enhance the reduced thrashing phenotype of *egl-4(ad450gf)* animals. Conversely, *npr-14(ok2375lf);egl-4(n479lf)* double mutants displayed high thrashing comparable to *egl-4(n479lf)* single mutants, rather than the reduced thrashing seen in *npr-14(ok2375lf)* animals (Figure 10b). Together, these results establish that NPR-14 acts upstream of EGL-4 in a linear pathway to regulate behavioural state: EGL-4 promotes quiescence, while NPR-14 functions upstream of EGL-4 to limit quiescence to promote wakefulness.

**Figure 10.**
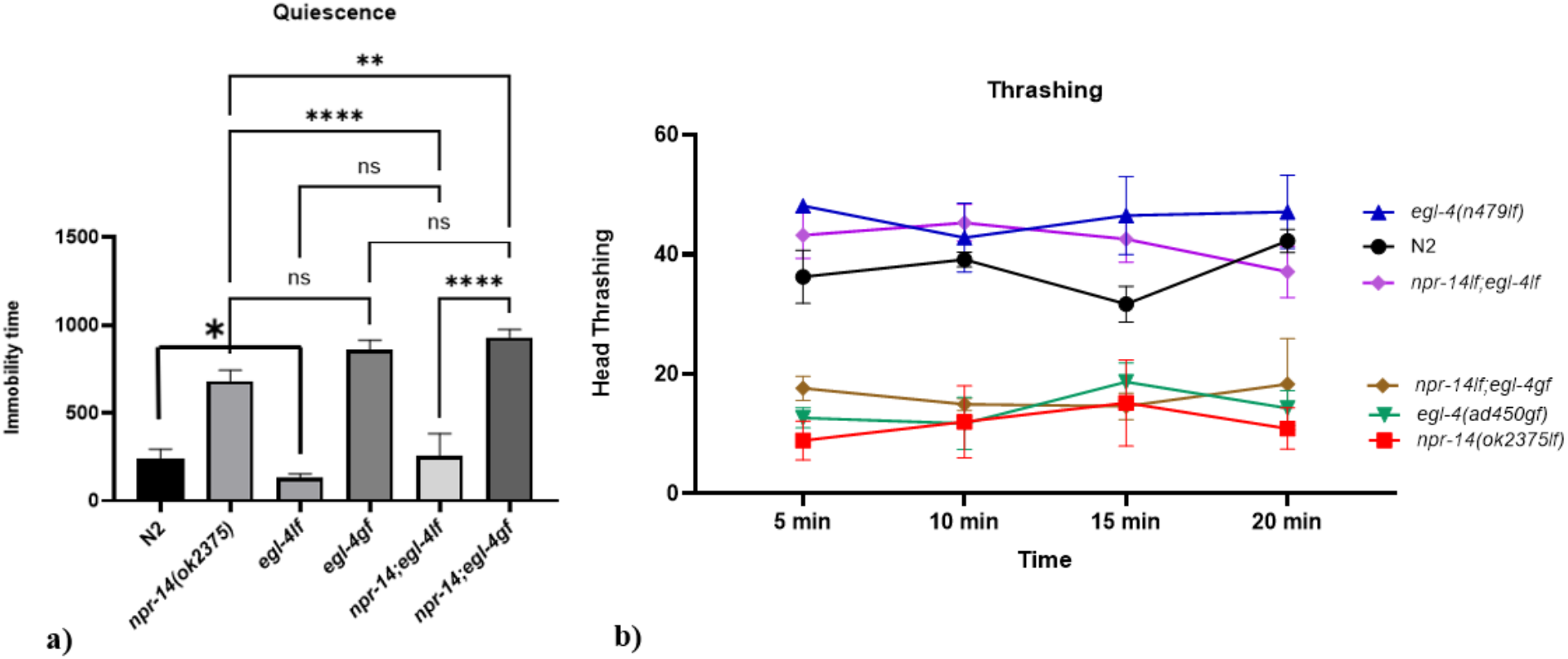
NPR-14 acts upstream of EGL-4 to regulate quiescence and locomotor activity. a) Total quiescence duration (seconds) over 30 minutes in wild-type N2, *npr-14(ok2375lf)*, *egl-4(lf)*, *egl-4(ad450gf)*, *npr-14(ok2375lf); egl-4(ad450gf)*, and *npr-14(ok2375lf);egl-4(n479lf)* animals. *npr-14(ok2375lf)* and *egl-4(ad450gf)* single mutants exhibit increased quiescence compared to wild-type. The *npr-14(ok2375lf);egl-4(ad450gf)* double mutant resembles *egl-4(ad450gf)* alone, while *npr-14(ok2375lf);egl-4(n479lf)* resembles *egl-4(n479lf)* alone, indicating that *egl-4* is epistatic to *npr-14*. b) Thrashing activity (head thrashes per 30 seconds) measured over 20 minutes. *npr-14(ok2375lf)* mutants show reduced thrashing compared to wild-type N2. *egl-4(n479lf)* mutants exhibit elevated thrashing, while *egl-4(ad450gf)* mutants show reduced thrashing. Double mutants display epistatic relationships consistent with panel (a): *npr-14(ok2375lf);egl-4(ad450gf)* resembles *egl-4(ad450gf)*, and *npr-14(ok2375lf);egl-4(n479lf)* resembles *egl-4(n479lf)*. Data are mean ± SEM; *p < 0.01, **p < 0.001, ***p < 0.0001, ns = not significant.

### Caffeine Selectively Rescues Quiescence in High-Sleep Genotypes

Because NPR-14 promotes arousal, similar to the wake-promoting function of orexin receptors in promoting wakefulness, we tested whether caffeine—a pharmacological arousal stimulus—could reverse the enhanced quiescence of *npr-14(ok2375lf)* mutants. In mammals, orexin/hypocretin signalling maintains wakefulness, while adenosine promotes sleep; caffeine antagonizes adenosine receptors to promote arousal (19,58,59). We examined whether caffeine could suppress the reduced-movement phenotype of *npr-14(ok2375lf)* mutants, revealing functional interactions between NPR-14-dependent quiescence and conserved arousal mechanisms. We measured thrashing activity in the presence or absence of caffeine to determine whether *npr-14* and *egl-4* regulate sensitivity to external arousal stimuli. Caffeine significantly increased thrashing in “sleepy” genotypes with elevated quiescence: *npr-14(ok2375lf)* single mutants, *egl-4(ad450gf)* single mutants, and *npr-14(ok2375lf);egl-4(ad450gf)* double mutants (Figure 11). In contrast, caffeine had minimal effect on wild-type N2 animals or on hyperactive “low-sleep” *egl-4(n479lf)* single mutants.

**Figure 11.**
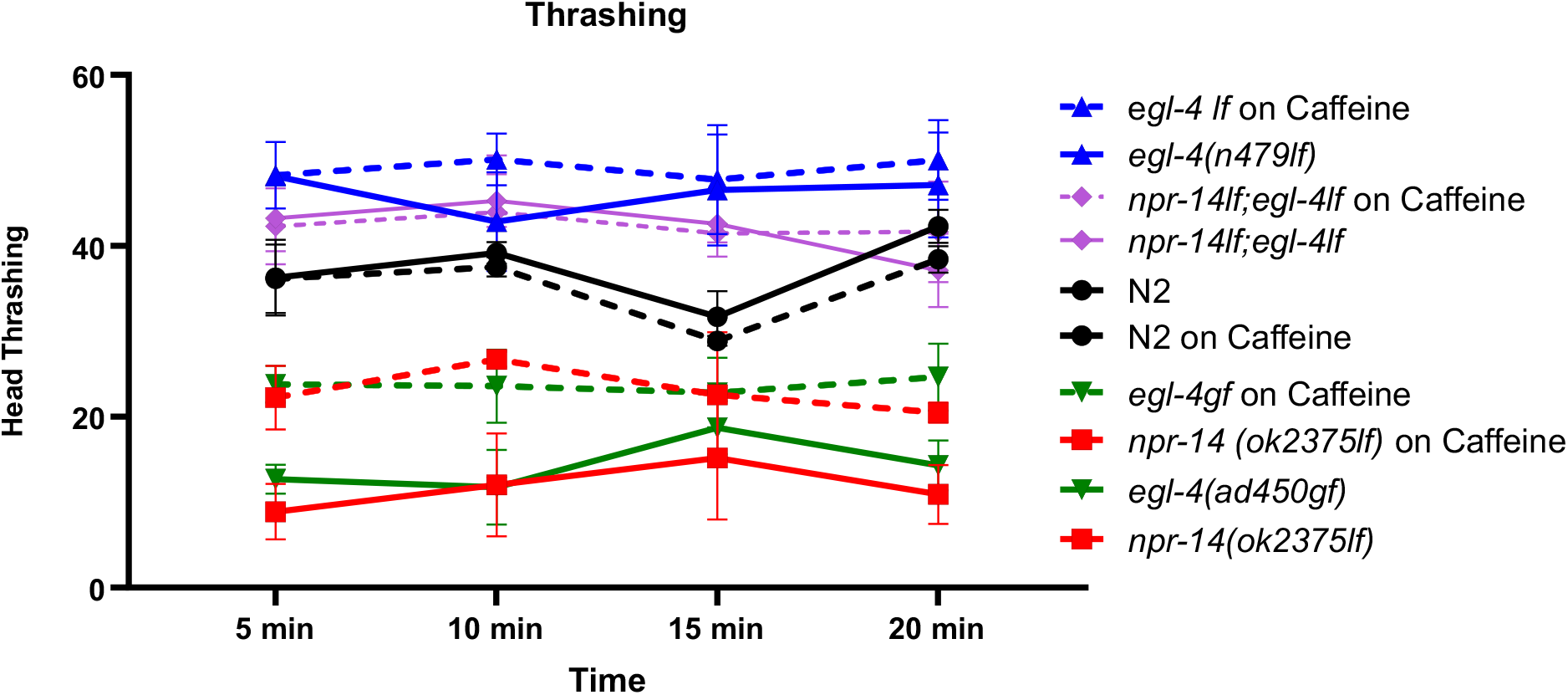
Caffeine selectively rescues quiescence in high-sleep genotypes. Thrashing activity (head thrashes per 30 seconds) measured over 20 minutes in M9 buffer alone (basal) or M9 supplemented with caffeine. Caffeine significantly increases thrashing in *npr-14(ok2375lf)*, *egl-4(ad450gf)*, and *npr-14(ok2375);egl-4(ad450gf)* mutants, but has minimal effect on wild-type N2, *egl-4(n479lf)*, or *npr-14(ok2375lf);egl-4(n479lf)* animals. Data are mean ± SEM; statistical significance indicated.

## Discussion

In this study, we identify the *C. elegans* neuropeptide receptor NPR-14 as a critical component of the arousal-promoting signaling pathway that constrains sleep-like quiescence. Our findings establish NPR-14 as an upstream inhibitor of the cGMP-dependent protein kinase EGL-4, defining a novel molecular axis for sleep-wake regulation.

### NPR-14 Functions as a Wake-Promoting Neuropeptide Receptor

The loss-of-function phenotypes observed in *npr-14* mutants—prolonged quiescence, reduced locomotion, impaired sensory responsiveness, decreased feeding and reproduction, and elevated lipid storage—collectively indicate a state of heightened sleep-like behavior. These phenotypes are rescued by transgenic expression of wild-type *npr-14*, confirming that they result specifically from loss of NPR-14 function. The expression of NPR-14 in ASH and ASI sensory neurons, along with GABAergic DD, VD, and VC motor neurons, positions this receptor to integrate environmental sensory input with motor output and metabolic state (49). This expression in various neurons suggests NPR-14 acts as a “gatekeeper” of arousal, preventing inappropriate entry into quiescence when environmental conditions demand active foraging and reproduction. The phenotypic similarity between *npr-14* loss-of-function mutants and gain-of-function mutations in *egl-4* (53) initially suggested these genes might function in opposition. Indeed, our epistasis analysis demonstrates that NPR-14 acts upstream of EGL-4 to regulate sleep-like quiescence. The observation that *egl-4(n479lf)* completely suppresses the quiescence of *npr-14(ok2375lf)* mutants, while *npr-14* loss does not enhance *egl-4(ad450gf)* phenotypes, indicates that NPR-14 normally functions to inhibit EGL-4 activity. This places NPR-14 as a wake-promoting receptor that constrains the sleep-promoting effects of PKG signaling.

### Mechanism of NPR-14 Action in the EGL-4 Pathway

EGL-4 promotes sleep-like quiescence by antagonizing cAMP-dependent signaling pathways (46,47). In mammals, PKG promotes sleep by opposing wake-promoting cAMP/PKA signaling, suggesting a conserved regulatory axis between PKG and cAMP signaling. The mechanism by which NPR-14 inhibits EGL-4 is currently unknown. As a GPCR, NPR-14 likely modulates cyclic nucleotide levels or calcium signaling through G-protein-mediated pathways. By analogy to mammalian orexin receptors, NPR-14 may couple to Gαq to activate phospholipase C and downstream calcium signaling, or potentially to Gαs to stimulate adenylyl cyclase and elevate cAMP, which could oppose EGL-4/PKG signaling at the pathway level (60,61). While our genetic data demonstrate that NPR-14 acts upstream of EGL-4, we cannot distinguish whether NPR-14 directly inhibits EGL-4 activity or acts in parallel to antagonize EGL-4-dependent outputs. Biochemical studies will be required to determine the molecular mechanism of this genetic interaction. Future studies examining the G-protein coupling specificity of NPR-14 and its effects on intracellular cAMP and cGMP levels will be essential to elucidate this mechanism.

NLP-59 has been proposed as a candidate orexin/allatotropin-like ligand for NPR-14 based on sequence homology (49), though this interaction has not been experimentally validated. RFamide peptides have been extensively implicated in the modulation of sleep-like states in *C. elegans*—FLP-13 and FLP-11 promote quiescence through receptors DMSR-1 and FRPR-4 (41,42,44,45), while NLP-22 regulates a sleep-like state through an unknown receptor (48). The identification of NPR-14 as a wake-promoting RFamide receptor establishes an opposing function for this peptide family in arousal regulation. Whether NLP-59 is the endogenous ligand for NPR-14, and whether additional ligands exist, remain important questions for future investigation.

### Integration with Conserved Sleep-Wake Circuitry

The NPR-14-EGL-4 pathway functions within a broader network of sleep-regulatory signals in *C. elegans*. NPR-14 joins a growing set of wake-promoting receptors in *C. elegans*, including NPR-9 which regulates local search behavior (62). Whether these receptors act in parallel or convergent pathways remains to be determined. Stress-induced sleep (SIS) is initiated by EGF signaling through LET-23/EGFR, which activates the sleep-promoting neurons ALA and RIS (36,39,40). These neurons release FLP-13 and FLP-11 neuropeptides that act via DMSR-1 and FRPR-4 receptors to suppress cAMP/PKA signaling and promote quiescence (46,47). NPR-14 likely acts in parallel or downstream of these pathways to modulate the probability of quiescence entry. The expression of NPR-14 in sensory neurons (ASH, ASI) that detect environmental cues suggests it may integrate sensory information with the sleep-promoting machinery to gate transitions between active and quiescent states.

Our finding that caffeine selectively rescues quiescence in high-sleep genotypes (*npr-14(lf)*, *egl-4(gf)*) but has minimal effect on wild-type or low-sleep *egl-4(lf)* animals suggests that NPR-14-dependent arousal pathways remain responsive to pharmacological stimulation. In mammals, caffeine antagonizes adenosine A1 receptors to promote wakefulness (19,24). While *C. elegans* lacks clear adenosine receptor orthologs, the observed caffeine sensitivity may reflect convergence on cAMP/PKA signaling or other conserved downstream effectors (59). Alternatively, caffeine may act through other mechanisms in worms, such as inhibition of phosphodiesterases or modulation of dopaminergic signaling (59). The selective rescue of high-quiescence genotypes suggests that NPR-14 and EGL-4 define a pathway that sets the threshold for arousal, which can be pharmacologically modulated when the system is biased toward sleep.

### Evolutionary Conservation of Arousal Mechanisms

NPR-14 belongs to the orexin/allatotropin receptor family, which includes vertebrate orexin/hypocretin receptors (OX1R, OX2R) that stabilize wakefulness by exciting monoaminergic systems (8,9,20). Loss of orexin signaling causes narcolepsy with cataplexy in humans and animal models (20). The sudden locomotor arrests observed in *npr-14(ok2375lf)* mutants superficially resemble narcoleptic episodes; however, whether this represents conservation of arousal mechanisms or convergent evolution of motor phenotypes remains unclear. Interestingly, the functional relationship between orexin/allatotropin receptors and PKG signaling appears to be evolutionarily conserved but with species-specific variations.

In *Drosophila*, high PKG levels increase wakefulness (33), opposite to the sleep-promoting role of EGL-4 in *C. elegans* (34). However, the fundamental antagonism between orexin signaling and sleep-promoting pathways is maintained—NPR-14 promotes wakefulness by inhibiting EGL-4, while in flies, the allatotropin receptor (when present) would presumably oppose sleep-promoting signals. The *Drosophila* allatotropin receptor has been lost, but the allatostatin-A receptor DAR-2 (phylogenetically related to galanin receptors) promotes sleep (27,63), suggesting that different GPCR families have evolved opposing roles in sleep regulation across species.

### Coupling of Behavioral and Metabolic States

The coordinated changes in locomotion, feeding, reproduction, and lipid storage observed in *npr-14* mutants reflect the tight coupling between sleep-like quiescence and metabolic regulation. In mammals, orexin neurons in the lateral hypothalamus not only promote wakefulness but also regulate feeding behavior and energy homeostasis (8,20). Similarly, NPR-14 appears to coordinate multiple aspects of physiology with behavioral state. The elevated fat accumulation in *npr-14* mutants may result from reduced energy expenditure during prolonged quiescence, or direct effects on metabolic regulation. The reduced egg-laying is consistent with the observation that reproductive behavior is suppressed during stress-induced sleep (55) and reflects the state-dependent coupling between locomotor output and reproduction. Because NPR-14 is expressed in VC motor neurons that directly control egg-laying by innervating vulval muscles (56,57), this receptor provides the physical connection needed to couple locomotor activity with reproductive output. This ensures both behaviors are suppressed during sleep-like quiescence.

## Conclusions and Future Directions

This study establishes NPR-14 as an arousal-promoting GPCR that inhibits EGL-4/PKG-dependent quiescence to maintain wakefulness. The NPR-14-EGL-4 axis represents a conserved molecular framework for sleep-wake regulation that links neuropeptidergic signaling to sensory-motor integration and metabolic homeostasis. Future studies should address: (1) the identity of the G-proteins coupled to NPR-14 and the downstream signaling mechanism; (2) the regulation of NLP-59 release and its role in modulating NPR-14 activity; (3) the interaction between NPR-14 and other sleep-regulatory neuropeptide systems; and (4) the molecular targets of EGL-4 that mediate quiescence downstream of NPR-14 inhibition. Understanding this pathway in *C. elegans* may provide insights into the molecular logic underlying sleep disorders such as narcolepsy and inform the development of novel therapeutic strategies for sleep regulation.

## References

1. Chaput JP, McNeil J, Després JP, Bouchard C, Tremblay A. Seven to Eight Hours of Sleep a Night Is Associated with a Lower Prevalence of the Metabolic Syndrome and Reduced Overall Cardiometabolic Risk in Adults. PLoS One. 2013 Sep 5;8(9):e72832. doi:10.1371/journal.pone.0072832 PubMed PMID: 24039808; PubMed Central PMCID: PMC3764138.

2. Chaput JP, McNeil J, Després JP, Bouchard C, Tremblay A. Short sleep duration as a risk factor for the development of the metabolic syndrome in adults. Preventive Medicine. 2013 Dec 1;57(6):872–7. doi:10.1016/j.ypmed.2013.09.022

3. Khan MA, Al-Jahdali H. The consequences of sleep deprivation on cognitive performance. Neurosciences (Riyadh). 2023 Apr;28(2):91–9. doi:10.17712/nsj.2023.2.20220108 PubMed PMID: 37045455; PubMed Central PMCID: PMC10155483.

4. Gottlieb DJ, Redline S, Nieto FJ, Baldwin CM, Newman AB, Resnick HE, et al. Association of usual sleep duration with hypertension: the Sleep Heart Health Study. Sleep. 2006 Aug;29(8):1009–14. doi:10.1093/sleep/29.8.1009 PubMed PMID: 16944668.

5. Knutson KL. Sleep duration and cardiometabolic risk: a review of the epidemiologic evidence. Best Pract Res Clin Endocrinol Metab. 2010 Oct;24(5):731–43. doi:10.1016/j.beem.2010.07.001 PubMed PMID: 21112022; PubMed Central PMCID: PMC3011978.

6. Garbarino S, Lanteri P, Bragazzi NL, Magnavita N, Scoditti E. Role of sleep deprivation in immune-related disease risk and outcomes. Commun Biol. 2021 Nov 18;4:1304. doi:10.1038/s42003-021-02825-4 PubMed PMID: 34795404; PubMed Central PMCID: PMC8602722.

7. Dijk D, Czeisler C. Contribution of the circadian pacemaker and the sleep homeostat to sleep propensity, sleep structure, electroencephalographic slow waves, and sleep spindle activity in humans. J Neurosci. 1995 May 1;15(5):3526–38. doi:10.1523/JNEUROSCI.15-05-03526.1995 PubMed PMID: 7751928; PubMed Central PMCID: PMC6578184.

8. Saper CB, Scammell TE, Lu J. Hypothalamic regulation of sleep and circadian rhythms. Nature. 2005 Oct;437(7063):1257–63. doi:10.1038/nature04284

9. Scammell TE, Arrigoni E, Lipton J. Neural Circuitry of Wakefulness and Sleep. Neuron. 2017 Feb 22;93(4):747–65. doi:10.1016/j.neuron.2017.01.014 PubMed PMID: 28231463; PubMed Central PMCID: PMC5325713.

10. Gompf HS, Anaclet C. The neuroanatomy and neurochemistry of sleep-wake control. Curr Opin Physiol. 2020 Jun;15:143–51. doi:10.1016/j.cophys.2019.12.012 PubMed PMID: 32647777; PubMed Central PMCID: PMC7347132.

11. Sulaman BA, Wang S, Tyan J, Eban-Rothschild A. Neuro-orchestration of sleep and wakefulness. Nat Neurosci. 2023 Feb;26(2):196–212. doi:10.1038/s41593-022-01236-w PubMed PMID: 36581730; PubMed Central PMCID: PMC12714371.

12. Eban-Rothschild A, Appelbaum L, de Lecea L. Neuronal Mechanisms for Sleep/Wake Regulation and Modulatory Drive. Neuropsychopharmacology. 2018 Apr;43(5):937–52. doi:10.1038/npp.2017.294 PubMed PMID: 29206811; PubMed Central PMCID: PMC5854814.

13. Kostin A, Alam MdA, Saevskiy A, Yang C, Golshani P, Alam MdN. Calcium Dynamics of the Ventrolateral Preoptic GABAergic Neurons during Spontaneous Sleep-Waking and in Response to Homeostatic Sleep Demands. Int J Mol Sci. 2023 May 5;24(9):8311. doi:10.3390/ijms24098311 PubMed PMID: 37176016; PubMed Central PMCID: PMC10179316.

14. Tossell K, Yu X, Giannos P, Anuncibay Soto B, Nollet M, Yustos R, et al. Somatostatin neurons in prefrontal cortex initiate sleep-preparatory behavior and sleep via the preoptic and lateral hypothalamus. Nat Neurosci. 2023;26(10):1805–19. doi:10.1038/s41593-023-01430-4 PubMed PMID: 37735497; PubMed Central PMCID: PMC10545541.

15. Arrigoni E, Fuller PM. The Sleep-Promoting Ventrolateral Preoptic Nucleus: What Have We Learned over the Past 25 Years? Int J Mol Sci. 2022 Mar 8;23(6):2905. doi:10.3390/ijms23062905 PubMed PMID: 35328326; PubMed Central PMCID: PMC8954377.

16. Funk CM, Peelman K, Bellesi M, Marshall W, Cirelli C, Tononi G. Role of Somatostatin-Positive Cortical Interneurons in the Generation of Sleep Slow Waves. J Neurosci. 2017 Sep 20;37(38):9132–48. doi:10.1523/JNEUROSCI.1303-17.2017 PubMed PMID: 28821651; PubMed Central PMCID: PMC5607463.

17. de Lecea L, Huerta R. Hypocretin (orexin) regulation of sleep-to-wake transitions. Front Pharmacol. 2014 Feb 12;5:16. doi:10.3389/fphar.2014.00016 PubMed PMID: 24575043; PubMed Central PMCID: PMC3921570.

18. Peyron C, Faraco J, Rogers W, Ripley B, Overeem S, Charnay Y, et al. A mutation in a case of early onset narcolepsy and a generalized absence of hypocretin peptides in human narcoleptic brains. Nat Med. 2000 Sep;6(9):991–7. doi:10.1038/79690 PubMed PMID: 10973318.

19. Reichert CF, Deboer T, Landolt H. Adenosine, caffeine, and sleep–wake regulation: state of the science and perspectives. J Sleep Res. 2022 Aug;31(4):e13597. doi:10.1111/jsr.13597 PubMed PMID: 35575450; PubMed Central PMCID: PMC9541543.

20. Jacobson LH, Hoyer D, de Lecea L. Hypocretins (orexins): The ultimate translational neuropeptides. J Intern Med. 2022 May;291(5):533–56. doi:10.1111/joim.13406 PubMed PMID: 35043499.

21. Porkka-Heiskanen T, Kalinchuk AV. Adenosine, energy metabolism and sleep homeostasis. Sleep Med Rev. 2011 Apr;15(2):123–35. doi:10.1016/j.smrv.2010.06.005 PubMed PMID: 20970361.

22. Huang L, Zhu W, Li N, Zhang B, Dai W, Li S, et al. Functions and mechanisms of adenosine and its receptors in sleep regulation. Sleep Med. 2024 Mar;115:210–7. doi:10.1016/j.sleep.2024.02.012 PubMed PMID: 38373361.

23. Ma WX, Yuan PC, Zhang H, Kong LX, Lazarus M, Qu WM, et al. Adenosine and P1 receptors: Key targets in the regulation of sleep, torpor, and hibernation. Front Pharmacol. 2023 Mar 10;14:1098976. doi:10.3389/fphar.2023.1098976 PubMed PMID: 36969831; PubMed Central PMCID: PMC10036772.

24. Gardiner C, Weakley J, Burke LM, Roach GD, Sargent C, Maniar N, et al. The effect of caffeine on subsequent sleep: A systematic review and meta-analysis. Sleep Medicine Reviews. 2023 Jun 1;69:101764. doi:10.1016/j.smrv.2023.101764

25. Abhilash L, Shafer OT. A two-process model of Drosophila sleep reveals an inter-dependence between circadian clock speed and the rate of sleep pressure decay. Sleep. 2023 Oct 31;47(2):zsad277. doi:10.1093/sleep/zsad277 PubMed PMID: 37930351; PubMed Central PMCID: PMC11275470.

26. Shaw PJ, Cirelli C, Greenspan RJ, Tononi G. Correlates of Sleep and Waking in Drosophila melanogaster. Science. 2000 Mar 10;287(5459):1834–7. doi:10.1126/science.287.5459.1834

27. Chen J, Reiher W, Hermann-Luibl C, Sellami A, Cognigni P, Kondo S, et al. Allatostatin A Signalling in Drosophila Regulates Feeding and Sleep and Is Modulated by PDF. PLoS Genet. 2016 Sep 30;12(9):e1006346. doi:10.1371/journal.pgen.1006346 PubMed PMID: 27689358; PubMed Central PMCID: PMC5045179.

28. Reinhard N, Schubert FK, Bertolini E, Hagedorn N, Manoli G, Sekiguchi M, et al. The Neuronal Circuit of the Dorsal Circadian Clock Neurons in Drosophila melanogaster. Front Physiol. 2022 Apr 29;13:886432. doi:10.3389/fphys.2022.886432 PubMed PMID: 35574472; PubMed Central PMCID: PMC9100938.

29. Nässel DR, Broeck JV. Insulin/IGF signaling in Drosophila and other insects: factors that regulate production, release and post-release action of the insulin-like peptides. Cell Mol Life Sci. 2015 Oct 15;73(2):271–90. doi:10.1007/s00018-015-2063-3 PubMed PMID: 26472340; PubMed Central PMCID: PMC11108470.

30. Shang Y, Donelson NC, Vecsey CG, Guo F, Rosbash M, Griffith LC. Short Neuropeptide F Is a Sleep-Promoting Inhibitory Modulator. Neuron. 2013 Oct 2;80(1):171–83. doi:10.1016/j.neuron.2013.07.029 PubMed PMID: 24094110.

31. Lismont E, Vleugels R, Marchal E, Badisco L, Van Wielendaele P, Lenaerts C, et al. Molecular cloning and characterization of the allatotropin precursor and receptor in the desert locust, Schistocerca gregaria. Front Neurosci. 2015 Mar 12;9. doi:10.3389/fnins.2015.00084

32. Alzugaray ME, Bruno MC, Villalobos Sambucaro MJ, Ronderos JR. The Evolutionary History of The Orexin/Allatotropin GPCR Family: from Placozoa and Cnidaria to Vertebrata. Sci Rep. 2019 Jul 15;9:10217. doi:10.1038/s41598-019-46712-9 PubMed PMID: 31308431; PubMed Central PMCID: PMC6629687.

33. Donlea J, Leahy A, Thimgan MS, Suzuki Y, Hughson BN, Sokolowski MB, et al. foraging alters resilience/vulnerability to sleep disruption and starvation in Drosophila. Proc Natl Acad Sci U S A. 2012 Feb 14;109(7):2613–8. doi:10.1073/pnas.1112623109 PubMed PMID: 22308351; PubMed Central PMCID: PMC3289360.

34. Raizen DM, Zimmerman JE, Maycock MH, Ta UD, You Y jai, Sundaram MV, et al. Lethargus is a Caenorhabditis elegans sleep-like state. Nature. 2008 Jan;451(7178):569–72. doi:10.1038/nature06535

35. Hill AJ, Mansfield R, Lopez JMNG, Raizen DM, Van Buskirk C. Cellular Stress Induces a Protective Sleep-like State in C. elegans. Current Biology. 2014 Oct 20;24(20):2399–405. doi:10.1016/j.cub.2014.08.040 PubMed PMID: 25264259.

36. Hill AJ, Robinson B, Jones JG, Sternberg PW, Buskirk CV. Sleep drive is coupled to tissue damage via shedding of Caenorhabditis elegans EGFR ligand SISS-1. Nature Communications. 2024 Dec 30;15:10886. doi:10.1038/s41467-024-55252-4 PubMed PMID: 39738055.

37. Iannacone MJ, Um P, Grubbs JI, Linden AM van der, Raizen DM. Quiescence Enhances Survival during Viral Infection in Caenorhabditis elegans. J Neurosci. 2024 Aug 28;44(35). doi:10.1523/JNEUROSCI.1700-22.2024 PubMed PMID: 39060176.

38. Cowen MH, Raizen DM, Hart MP. Structural neuroplasticity after sleep loss modifies behavior and requires neurexin and neuroligin. iScience. 2024 Apr 19;27(4):109477. doi:10.1016/j.isci.2024.109477 PubMed PMID: 38551003; PubMed Central PMCID: PMC10973677.

39. van Buskirk C, Sternberg PW. Epidermal growth factor signaling induces behavioral quiescence in Caenorhabditis elegans. Nature Neuroscience. 2007 Oct 1;10(10):1300–7. Located at: 26854954. doi:10.1038/nn1981

40. Nath RD, Chow ES, Wang H, Schwarz EM, Sternberg PW. C. elegans stress-induced sleep emerges from the collective action of multiple neuropeptides. Curr Biol. 2016 Sep 26;26(18):2446–55. doi:10.1016/j.cub.2016.07.048 PubMed PMID: 27546573; PubMed Central PMCID: PMC5694219.

41. Nelson MD, Lee KH, Churgin MA, Hill AJ, Van Buskirk C, Fang-Yen C, et al. FMRFamide-like FLP-13 neuropeptides promote quiescence following heat stress in Caenorhabditis elegans. Curr Biol. 2014 Oct 20;24(20):2406–10. doi:10.1016/j.cub.2014.08.037 PubMed PMID: 25264253; PubMed Central PMCID: PMC4254296.

42. Iannacone MJ, Beets I, Lopes LE, Churgin MA, Fang-Yen C, Nelson MD, et al. The RFamide receptor DMSR-1 regulates stress-induced sleep in C. elegans. eLife. 6:e19837. doi:10.7554/eLife.19837 PubMed PMID: 28094002; PubMed Central PMCID: PMC5241116.

43. Trojanowski NF, Raizen DM. Call it Worm Sleep. Trends in Neurosciences. 2016 Feb 1;39(2):54–62. doi:10.1016/j.tins.2015.12.005

44. Turek M, Besseling J, Spies JP, König S, Bringmann H. Sleep-active neuron specification and sleep induction require FLP-11 neuropeptides to systemically induce sleep. eLife. 5:e12499. doi:10.7554/eLife.12499 PubMed PMID: 26949257; PubMed Central PMCID: PMC4805538.

45. Rossi L, Amoako K, Busack I, Golinelli L, Courtney A, Besseling J, et al. The neuropeptide FLP-11 induces and self-inhibits sleep through the receptor DMSR-1 in Caenorhabditis elegans. Current Biology. 2025 May 5;35(9):2183–2194.e10. doi:10.1016/j.cub.2025.03.039 PubMed PMID: 40273913.

46. Trojanowski NF, Nelson MD, Flavell SW, Fang-Yen C, Raizen DM. Distinct Mechanisms Underlie Quiescence during Two Caenorhabditis elegans Sleep-Like States. J Neurosci. 2015 Oct 28;35(43):14571–84. doi:10.1523/JNEUROSCI.1369-15.2015 PubMed PMID: 26511247; PubMed Central PMCID: PMC4623228.

47. Cianciulli A, Yoslov L, Buscemi K, Sullivan N, Vance RT, Janton F, et al. Interneurons Regulate Locomotion Quiescence via Cyclic Adenosine Monophosphate Signaling During Stress-Induced Sleep in Caenorhabditis elegans. Genetics. 2019 Sep;213(1):267–79. doi:10.1534/genetics.119.302293 PubMed PMID: 31292211; PubMed Central PMCID: PMC6727807.

48. Nelson M, Trojanowski N, George-Raizen J, Smith C, Yu CC, Fang-Yen C, et al. The neuropeptide NLP-22 regulates a sleep-like state in Caenorhabditis elegans. Nat Commun. 2013;4:2846. doi:10.1038/ncomms3846 PubMed PMID: 24301180; PubMed Central PMCID: PMC3867200.

49. Beets I, Zels S, Vandewyer E, Demeulemeester J, Caers J, Baytemur E, et al. System-wide mapping of peptide-GPCR interactions in C. elegans. Cell Rep. 2023 Sep 26;42(9):113058. doi:10.1016/j.celrep.2023.113058 PubMed PMID: 37656621; PubMed Central PMCID: PMC7615250.

50. Waterhouse AM, Procter JB, Martin DMA, Clamp M, Barton GJ. Jalview Version 2—a multiple sequence alignment editor and analysis workbench. Bioinformatics. 2009 May 1;25(9):1189–91. doi:10.1093/bioinformatics/btp033 PubMed PMID: 19151095; PubMed Central PMCID: PMC2672624.

51. López-Cruz A, Sordillo A, Pokala N, Liu Q, McGrath PT, Bargmann CI. Parallel Multimodal Circuits Control an Innate Foraging Behavior. Neuron. 2019 Apr 17;102(2):407–419.e8. doi:10.1016/j.neuron.2019.01.053 PubMed PMID: 30824353.

52. Mclntire SL, Jorgensen E, Kaplan J, Horvitz HR. The GABAergic nervous system of Caenorhabditis elegans. Nature. 1993 Jul;364(6435):337–41. doi:10.1038/364337a0

53. Raizen DM, Cullison KM, Pack AI, Sundaram MV. A Novel Gain-of-Function Mutant of the Cyclic GMP-Dependent Protein Kinase egl-4 Affects Multiple Physiological Processes in Caenorhabditis elegans. Genetics. 2006 May;173(1):177–87. doi:10.1534/genetics.106.057380 PubMed PMID: 16547093; PubMed Central PMCID: PMC1461420.

54. Daniels SA, Ailion M, Thomas JH, Sengupta P. egl-4 acts through a transforming growth factor-beta/SMAD pathway in Caenorhabditis elegans to regulate multiple neuronal circuits in response to sensory cues. Genetics. 2000 Sep;156(1):123–41. doi:10.1093/genetics/156.1.123 PubMed PMID: 10978280; PubMed Central PMCID: PMC1461244.

55. Subramanian S, Diya N, Nelson MD. Egg laying during stress-induced sleep of Caenorhabditis elegans is reduced due to behavioral quiescence and fertility defects. MicroPubl Biol. 2025:10.17912/micropub.biology.001735. doi:10.17912/micropub.biology.001735 PubMed PMID: 40741294; PubMed Central PMCID: PMC12308320.

56. Kopchock RJ, Ravi B, Bode A, Collins KM. The Sex-Specific VC Neurons Are Mechanically Activated Motor Neurons That Facilitate Serotonin-Induced Egg Laying in C. elegans. J Neurosci. 2021 Apr 21;41(16):3635–50. doi:10.1523/JNEUROSCI.2150-20.2021 PubMed PMID: 33687965; PubMed Central PMCID: PMC8055074.

57. Zhang M, Chung SH, Fang-Yen C, Craig C, Kerr RA, Suzuki H, et al. A Self-Regulating Feed-Forward Circuit Controlling C. elegans Egg-Laying Behavior. Current Biology. 2008 Oct 14;18(19):1445–55. doi:10.1016/j.cub.2008.08.047 PubMed PMID: 18818084.

58. Aldosari MS, Olaish AH, Nashwan SZ, Abulmeaty MMA, BaHammam AS. The effects of caffeine on drowsiness in patients with narcolepsy: a double-blind randomized controlled pilot study. Sleep Breath. 2020 Dec;24(4):1675–84. doi:10.1007/s11325-020-02065-6 PubMed PMID: 32215834.

59. Bridi JC, Barros AG de A, Sampaio LR, Ferreira JCD, Antunes Soares FA, Romano-Silva MA. Lifespan Extension Induced by Caffeine in Caenorhabditis elegans is Partially Dependent on Adenosine Signaling. Front Aging Neurosci. 2015 Dec 8;7:220. doi:10.3389/fnagi.2015.00220 PubMed PMID: 26696878; PubMed Central PMCID: PMC4672644.

60. Holmqvist T, Johansson L, Östman M, Ammoun S, Åkerman KEO, Kukkonen JP. OX1 Orexin Receptors Couple to Adenylyl Cyclase Regulation via Multiple Mechanisms *. Journal of Biological Chemistry. 2005 Feb 25;280(8):6570–9. doi:10.1074/jbc.M407397200 PubMed PMID: 15611118.

61. Lund PE, Shariatmadari R, Uustare A, Detheux M, Parmentier M, Kukkonen JP, et al. The Orexin OX1 Receptor Activates a Novel Ca2+ Influx Pathway Necessary for Coupling to Phospholipase C *. Journal of Biological Chemistry. 2000 Oct 6;275(40):30806–12. doi:10.1074/jbc.M002603200 PubMed PMID: 10880509.

62. Bendena WG, Boudreau JR, Papanicolaou T, Maltby M, Tobe SS, Chin-Sang ID. A Caenorhabditis elegans allatostatin/galanin-like receptor NPR-9 inhibits local search behavior in response to feeding cues. Proceedings of the National Academy of Sciences. 2008 Jan 29;105(4):1339–42. doi:10.1073/pnas.0709492105

63. Wegener C, Chen J. Allatostatin A Signalling: Progress and New Challenges From a Paradigmatic Pleiotropic Invertebrate Neuropeptide Family. Front Physiol. 2022 Jun 24;13:920529. doi:10.3389/fphys.2022.920529 PubMed PMID: 35812311; PubMed Central PMCID: PMC9263205.

64. Brenner S. The Genetics of CAENORHABDITIS ELEGANS. Genetics. 1974 May;77(1):71–94. doi:10.1093/genetics/77.1.71 PubMed PMID: 4366476; PubMed Central PMCID: PMC1213120.

65. Rieckher M, Tavernarakis N. Generation of Caenorhabditis elegans Transgenic Animals by DNA Microinjection. Bio Protoc. 2017 Oct 5;7(19):e2565. doi:10.21769/BioProtoc.2565 PubMed PMID: 29071286; PubMed Central PMCID: PMC5734614.

66. Hobert O. PCR fusion-based approach to create reporter gene constructs for expression analysis in transgenic C. elegans. Biotechniques. 2002 Apr;32(4):728–30. doi:10.2144/02324bm01 PubMed PMID: 11962590.

67. Perkins LA, Hedgecock EM, Thomson JN, Culotti JG. Mutant sensory cilia in the nematode *Caenorhabditis elegans*. Developmental Biology. 1986 Oct 1;117(2):456–87. doi:10.1016/0012-1606(86)90314-3

68. Wang FY, Ching TT. Oil Red O Staining for Lipid Content in Caenorhabditis elegans. Bio Protoc. 2021 Aug 20;11(16):e4124. doi:10.21769/BioProtoc.4124 PubMed PMID: 34541042; PubMed Central PMCID: PMC8413582.

69. Min H, Youn E, Shim YH. Long-Term Caffeine Intake Exerts Protective Effects on Intestinal Aging by Regulating Vitellogenesis and Mitochondrial Function in an Aged Caenorhabditis Elegans Model. Nutrients. 2021 Jul 23;13(8):2517. doi:10.3390/nu13082517 PubMed PMID: 34444677; PubMed Central PMCID: PMC8398797.

